# Motor learning drives region-specific transcriptomic remodeling in the motor cortex and dorsal striatum

**DOI:** 10.1101/2025.07.11.664268

**Authors:** Yue Sun, Richard H. Roth, Fuu-Jiun Hwang, Sui Wang, Jun B. Ding

## Abstract

Motor learning depends on coordinated activity across the motor cortex (M1) and dorsal striatum (dSTR), yet the molecular mechanisms driving learning-related synaptic and circuit remodeling remain unclear. Here, we combine activity-dependent genetic labeling (TRAP) with single-cell RNA sequencing to generate an unbiased, cell type-resolved transcriptional atlas of behaviorally engaged populations during a forelimb reaching task. We identify diverse activated neurons across M1 and dSTR, including a striking enrichment of Htr3a-expressing interneurons (Htr3a INs) in M1 that are selectively recruited during skilled reaching, as confirmed by two-photon calcium imaging. Corticostriatal projection neurons and striatal spiny projection neurons show subtype-and region-specific transcriptional remodeling involving genes linked to synaptic function, translation, and metabolism. Glial cells—including astrocytes, oligodendrocytes, and microglia— exhibit similarly robust, stage-and region-dependent gene regulation. These findings provide a comprehensive molecular framework for motor learning and highlight coordinated, cell type-specific transcriptional programs in neurons and glia that shape the encoding and retrieval of motor memory.

**Highlights:**

- Motor learning activates interneuron cell types in motor cortex and striatum
- Htr3a-expressing interneurons in motor cortex are specifically activated while performing a learned reaching behavior
- Transcriptome remodeling exhibited distinct patterns between motor cortex and striatum
- Glial cells showed stage- and region-specific transcriptomic alteration patterns that align with those in neurons

## Introduction

The ability to acquire and execute skilled movements is a fundamental feature of animal behavior and is central to our interactions with the external world. These abilities are shaped by motor learning, a process through which repeated practice leads to the formation of long-lasting motor memories and the refinement of motor performance. At the circuit level, the projection from the primary motor cortex (M1) to the dorsal striatum (dSTR) serves as a key substrate for both the acquisition and retrieval of learned motor behaviors. This corticostriatal pathway integrates planning, execution, and evaluation of movement and is essential for encoding the temporal and spatial precision required for skilled actions. Disruptions to this projection impair motor skill learning and compromise the recall of previously acquired movements, highlighting its essential role in the formation and expression of motor memory^1^.

Despite a wealth of behavioral and physiological studies that have characterized learning-related changes in this circuit, the molecular mechanisms that govern the formation, consolidation, and retrieval of motor memories remain incompletely understood. *In vivo* calcium imaging, electrophysiological recordings, and structural studies have revealed extensive functional and anatomical remodeling within both M1 and dSTR during motor learning. For example, excitatory projection neurons in M1 layer 2/3 (L2/3) and layer 5 (L5) show learning-dependent tuning to reach trajectories^2,3^, while spiny projection neurons (SPNs) in dSTR develop task-locked firing patterns that mirror learned movements^4^. Structural plasticity, including dendritic spine formation and elimination in M1^5–9^ and bouton remodeling in inhibitory interneurons^10^, further supports the idea that motor learning induces profound circuit-level rewiring. However, these observations, while compelling, have largely outpaced our understanding of the underlying transcriptional programs that enable these plastic changes. In particular, the molecular events that support the encoding of new motor skills and the retrieval of consolidated motor memories remain poorly defined at the level of specific cell types within this circuit.

A comprehensive understanding of motor learning and memory requires a shift from focusing solely on electrophysiological changes or individual molecular pathways to embracing an unbiased, system-level approach capable of resolving dynamic gene expression programs across the full cellular diversity of the motor circuit. This is especially important given the heterogeneity of cell types that contribute to learning-related plasticity. Inhibitory interneurons—including somatostatin (SST)-, parvalbumin (PV)-, and vasoactive intestinal peptide (VIP)-expressing populations—exhibit plasticity in firing patterns and connectivity during motor training^11–14^. These interneurons shape local computation and are increasingly recognized as integral components of motor memory ensembles^10,15^. Glial cells are also emerging as critical regulators of synaptic and circuit-level plasticity. Astrocytes influence neurotransmission through tripartite synapses and have been shown to modulate the formation of learning-associated neural assemblies^16,17^. Oligodendrocytes promote myelin remodeling in an activity-dependent manner that supports neural signal propagation and skill refinement^18–21^. Microglia dynamically sculpt synaptic connections *via* spine elimination during motor learning, contributing to long-term circuit refinement ^22,23^. Despite these emerging insights, the full transcriptional landscape of learning-induced plasticity across all major cell types and the coordination between them remains largely unexplored.

To address these gaps, we implemented an integrative, unbiased approach by combining the activity-dependent genetic labeling method TRAP (Targeted Recombination in Active Populations)^24^ with single-cell RNA sequencing (scRNA-seq) to profile transcriptional responses in neurons and glia within the motor cortex and dorsal striatum. We employed a single-pellet forelimb reaching task, a well-established paradigm for assessing skilled motor learning in mice, and captured cells active during both early learning and late, well-trained stages. This strategy enabled us to generate a cell type-resolved molecular atlas of motor learning, capturing both presynaptic M1 projection neurons and their postsynaptic targets in the striatum, as well as the surrounding interneuronal and glial landscape.

Our dataset reveals several key findings. First, we identify widespread transcriptional remodeling in projection neurons and interneurons, including a striking and selective recruitment of Htr3a-expressing interneurons (Htr3a INs) in the motor cortex during trained reaching. Two-photon calcium imaging confirmed their preferential activation during skilled movement execution. Second, we uncover robust, cell type-specific gene expression changes across astrocytes, oligodendrocytes, and microglia, revealing glial engagement that is both region-and learning stage-specific. Third, we show that synaptic, cytoskeletal, and signaling pathways are differentially regulated across M1 and dSTR, suggesting that distinct molecular strategies are employed across brain regions and cell types to support motor learning and memory consolidation.

Altogether, this work provides a high-resolution, unbiased transcriptional framework for understanding the molecular logic of motor learning across the corticostriatal circuit. By capturing activity-defined cellular states and identifying both shared and region-specific gene expression programs, our findings offer a powerful resource for dissecting the cellular and molecular mechanisms that enable the transformation of experience into lasting motor memory.

## Results

### Labeling cellular populations engaged during motor learning

To investigate cell type specific transcriptomic dynamics associated with motor learning and memory formation, we utilized the Targeted Recombination in Active Populations (TRAP) system^24^ to label behaviorally activated neurons in mice trained on a forelimb single-pellet reaching task. Following TRAP labeling, single-cell RNA sequencing (scRNA-seq) was performed on cells dissociated from the primary motor cortex (M1) and dorsal striatum (dSTR) (Figure 1A-1C). In TRAP mice, Cre^ER^ expression is driven by immediate-early gene promoters (*c-Fos* or *Arc*) and activated by 4-hydroxytamoxifen (4-OHT), enabling temporally controlled genetic labeling of neurons responsive to behavioral experience. To tag these active neurons with fluorescence, we crossed TRAP lines (Fos-TRAP and Arc-TRAP) with Cre-dependent tdTomato reporter mice (Ai14 or Ai9). Mice were trained to reach, grasp, and retrieve food pellets through a narrow slit using a single forelimb (Figure 1A). Their performance improved over eight daily training sessions, as measured by the success rate and the frequency of successful reaches (Figure 1D and see details in Methods). To label learning stage-specific neuronal ensembles, 4-OHT was administrated either during the first two daily training sessions (early TRAP) or the last two sessions after animals became proficient (late TRAP), resulting in permanent tdTomato expression in activated neurons (TRAP-tdTom). Early TRAP mice underwent only two days of training, while late TRAP mice completed all eight days of training. Control TRAP mice were exposed to the same training environment and received 4-OHT at the same time points as either early or late TRAP groups, but were not trained to perform the reaching task (Figure 1B). Due to known differences in TRAP efficiency between *c-Fos* and *Arc* drivers in cortex and striatum^24^, we used Fos-TRAP to label cortical neurons, while Arc-TRAP to label striatal neurons (Figure 1E, S1D and S1E). Tissues were collected three days after TRAP labeling for scRNA-seq from M1 in Fos-TRAP mice and dorsal striatum in Arc-TRAP mice.

**Figure 1.**
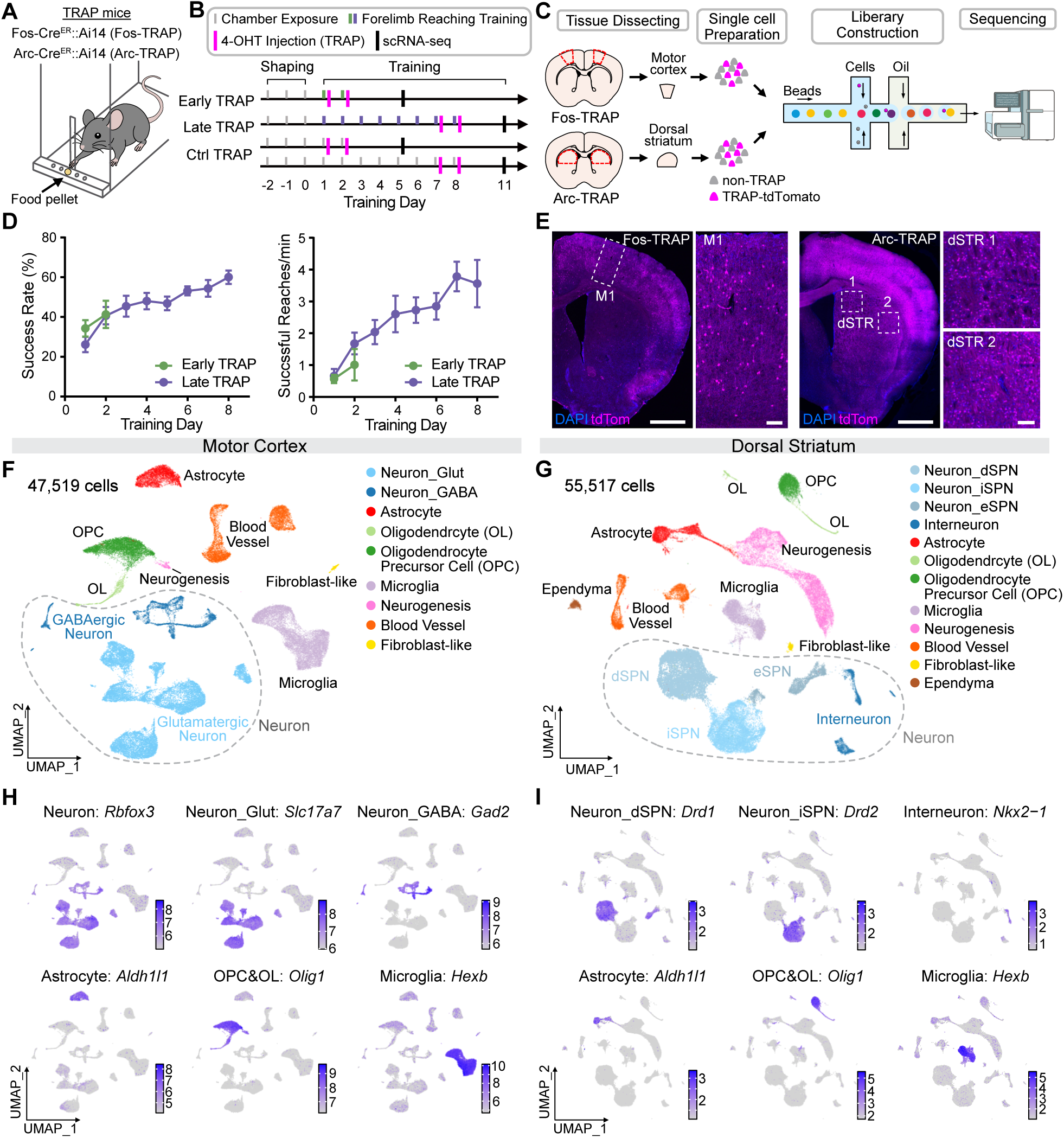
Single-cell transcriptomic profiling of behaviorally relevant cells in motor learning. (A) Schematic of the freely moving forelimb single-pellet reaching task. (B) Experimental timeline illustrating the reaching task and TRAP labeling schedule. (C) Overview of the single-cell RNA sequencing (scRNA-seq) workflow. (D) Behavioral performance of mice used for scRNA-seq, quantified by success rate (left) and successful reaches per minute (right). n= 8 Early TRAP (4 Fos-TRAP, 4 Arc-TRAP), n= 8 Late TRAP (4 Fos-TRAP, 4 Arc-TRAP). Error bars represent mean ± SEM. (E) Representative images showing TRAP-tdTomato+ cells (magenta) in Fos-TRAP (left) and Arc-TRAP (right) mice. Dashed squares indicate magnified regions of M1 and dorsal striatum (dSTR). Scale bars: whole brain, 1 mm; M1 and dSTRs, 50 μm. (F) Unbiased clustering and cell type annotation of 47,519 cells from motor cortex visualized using Uniform Manifold Approximation and Projection (UMAP). Colors denote major neuronal and glial cell types. (G) Unbiased clustering and cell type annotation of 55,517 dorsal striatum cells visualized by UMAP. Colors represent major neuronal and glial cell types. (H) Relative expression levels of marker genes for major neuronal and glial cell types across clusters in the motor cortex. (I) Relative expression of marker genes for major neuronal and glial cell types across clusters in the dorsal striatum. See also Figure S1 ans S2 Sun et al., *BioRxiv.* 2025 | Page 11

Previous work showed that Fos-TRAP labeling in M1 captures neurons activated during learning and reactivated during motor memory retrieval^9^. To assess whether Arc-TRAP labels motor memory-relevant neurons in the dSTR, we trained separate cohorts of Fos-TRAP and Arc-TRAP mice in a memory retrieval session and performed histological analyses (Figure S1A and S1B). TRAP-labeled neurons (TRAP-tdTom) represent cells activated during learning, while memory retrieval-activated neurons were identified by c-Fos immunostaining or *Arc* RNA *in situ* hybridization (ISH) (Figure S1B-S1D). Consistent with our previous finding in the motor cortex ^9^ , although the total number of TRAP-tdTom neurons was comparable across groups, the late TRAP group exhibited a significantly higher proportion of reactivated neurons (TRAP-tdTom and c-Fos double-positive) (Figure S1E), suggesting the emergence of memory-relevant ensembles over the course of learning. In contrast, while the number of TRAP-tdTom neurons in dSTR was significantly elevated in the Late TRAP group relative to controls, there was no corresponding increase in reactivation during memory retrieval (Figure S1F). These results suggest that the striatum, unlike the motor cortex, recruits more neurons as learning progresses but may not form stable reactivated ensembles associated with memory retrieval^25^. Altogether, these findings validate TRAP as an effective tool to label neurons involved in motor learning across both regions, while also highlighting distinct circuit dynamics in the motor cortex and striatum.

### Molecular profile of neuronal subtypes engaged during motor learning

To profile the learning-associated transcriptional changes in a cell type-specific manner, we performed scRNA-seq and used the detection of tdTomato transcripts as a proxy for TRAP-labeled, behaviorally activated neurons. Following quality control (Figure S2A and S2B; See details in Methods), we recovered 47,519 cells from the motor cortex and 55,517 cells from dorsal striatum and performed unbiased clustering using Seurat v4^26^ (Figure 1F, 1G, S2C and S2D). Fifty distinct clusters in motor cortex (Figure S2C) and 43 in dorsal striatum (Figure S2D) were identified and annotated based on their expression of known marker genes (Figure 1H,1I, S2E and S2F).

In the motor cortex, 23 neuronal clusters (20,344 neurons) were further categorized into ten glutamatergic projection neuron subtypes and nine GABAergic interneuron subtypes (Figure 2A and S3A). These subtypes were present in similar proportions across the control, early, and late TRAP groups, indicating that motor learning does not significantly alter cell composition (Figure 2B). Similarly, in dorsal striatum, 18 neuronal clusters (25,002 neurons), including three spiny projection neuron (SPN) subtypes and four interneuron subtypes (Figure 2C and S3B), also showed stable subtype distributions across learning stages (Figure 2D). The identified cell types and their marker gene expression profile (Figure S3C and S3D) were highly consistent with previously reported scRNA-seq atlases of the motor cortex^27–32^ and striatum^31,33,34^, validating the fidelity of our dataset.

**Figure 2.**
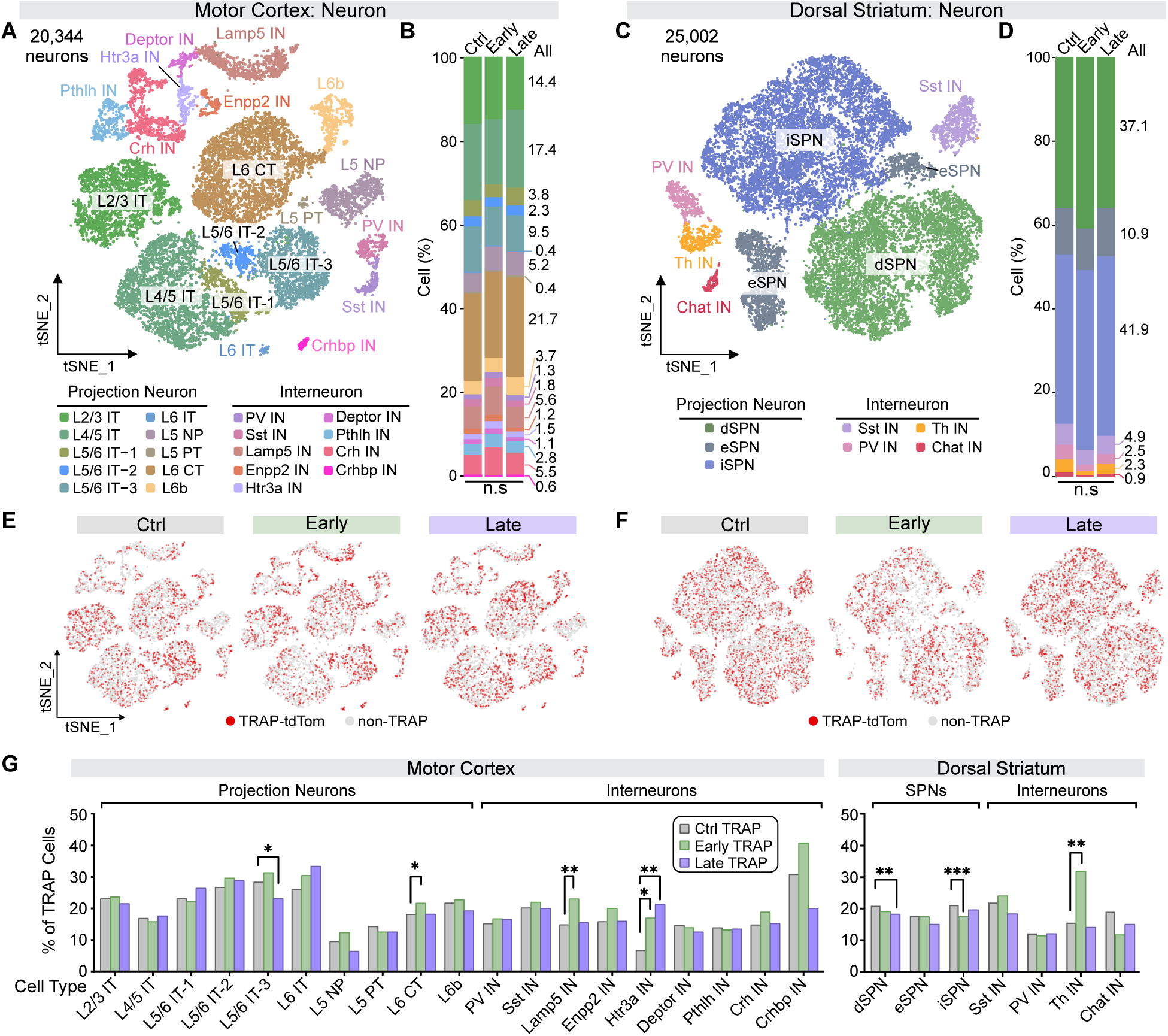
Motor learning activates diverse neuronal subtypes in the motor cortex and dorsal striatum. (A) Neuronal subtypes identified in the motor cortex. Top: t-distributed stochastic neighbor embedding (t-SNE) plots showing clustering of neuronal subtypes. Bottom: Corresponding neuronal subtype labels. Colors denote distinct neuronal subtypes. (B) Proportions of neuronal subtypes across Ctrl, Early, and Late TRAP groups in the motor cortex. Two-way ANOVA; ns, non-significant. (C) Neuronal subtypes identified in the dorsal striatum. Top: t-SNE plot of neuronal clusters. Bottom: Corresponding neuronal subtype labels. Colors represent distinct neuronal subtypes. (D) Proportions of neuronal subtypes across Ctrl, Early, and Late TRAP groups in the dorsal striatum. Two-way ANOVA; ns, non-significant. (E) Distribution of TRAP-tdTomato cells (highlight in red) across Ctrl, Early and Late TRAP groups in the motor cortex, visualized using t-SNE. (F) Distribution of TRAP-tdTomato cells (highlight in red) across Ctrl, Early and Late TRAP groups in the dorsal striatum, visualized using t-SNE. (G) Proportion of TRAP cells within each neuronal subtype across Ctrl, Early and Late TRAP groups in the motor cortex (left) and dorsal striatum (right). Bars represent proportions. Fisher’s exact test was used to compare TRAP and non-TRAP cells between the Ctrl and Early TRAP , and between the Ctrl and Late TRAP groups. *p < 0.05; **p < 0.01. Abbreviations: IT, intratelencephalically projecting; NP, near-projecting; PT, pyramidal tract; CT, corticothalamic projecting; PV, parvalbumin; SPN, spiny projection neuron. See also Figure S3 Sun et al., *BioRxiv.* 2025 | Page 13

We next assessed the distribution of TRAP-labeled (TRAP-tdTom) neurons across the neuronal subtypes (Figure S3E). We found that TRAP-tdTom neurons were broadly distributed across all neuronal types in both regions (Figure 2E, 2F and S3F). To identify cell types preferentially engaged during learning, we quantified the proportion of TRAP neurons in each neuronal subtype across control, early, and late TRAP groups (Figure 2G). In motor cortical projection neurons, although TRAP proportions were generally higher compared to interneurons, most subtypes did not show significant differences across learning stages. In M1, corticothalamic projection neurons in layer 6 (L6 CT) showed a significant increase in labeling in early TRAP, while one of the intratelencephalic projection neurons in layer 5 and 6 (L5/6 IT-3) showed a significant decrease in labeling in the late stage (Figure 2G). This suggests that during early learning, there is an expansion of the activated cortical ensemble, which is later followed by decreased activation and refinement of this ensemble in the late stage, consistent with previous functional measures ^2,35,36^. In the dorsal striatum, the fraction of TRAP direct pathway SPNs (dSPNs) decreased during late learning, while indirect pathway SPNs (iSPNs) showed reduced labeling in early TRAP (Figure 2G).

In contrast, several interneuron populations displayed marked learning stage-specific enrichment (Figure 2G). Cortical interneurons expressing lysosomal-associated membrane protein family member 5 (Lamp5 INs) were significantly enriched for TRAP cells during early learning, while 5-Hydroxytryptamine receptor 3a (Htr3a)-expressing interneurons (Htr3a INs) showed increased activation at both early and late stages. In the dorsal striatum, tyrosine hydroxylase-expressing interneurons (Th INs) were preferentially engaged during early learning. These findings point to a key, yet underexplored, role for specific interneuron subtypes in shaping the neural dynamics of motor learning. While their precise functional contributions remain to be defined, these cell types may participate in circuit-level plasticity underlying motor skill acquisition and memory consolidation.

### Activation of Htr3a-expressing interneurons is specific to learned motor behavior

Htr3a-expressing interneurons , along with parvalbumin (PV)-and somatostatin (SST)-expressing interneurons (INs), constitute the major principal classes of cortical interneurons ^37^. While distinct from PV INs and SST INs, recent single-cell transcriptomic studies have revealed that Htr3a INs represent a subtype of vasoactive intestinal peptide (VIP)-expressing interneurons (DropViz^31^; Cell Type Knowledge Explorer^32^). In our dataset, we confirmed that Htr3a INs co-express *Vip* but not *Sst* or *Pvalb*, comprising ∼13.5% of all VIP interneurons in the motor cortex (Figure 3A, S3C and S4A). Htr3a INs also express canonical VIP IN marker genes, including Cholecystokinin (*Cck*) and Neuropeptide Y (*Npy*), but lack expression of calretinin (*Calb2*) or Choline acetyltransferase (*Chat*) (Figure 3B, 3C and S4A). Notably, nearly all Htr3a INs expressed *Cck* (99.7%; Figure 3C), defining them as a subtype of Cck^+^ VIP INs.

**Figure 3.**
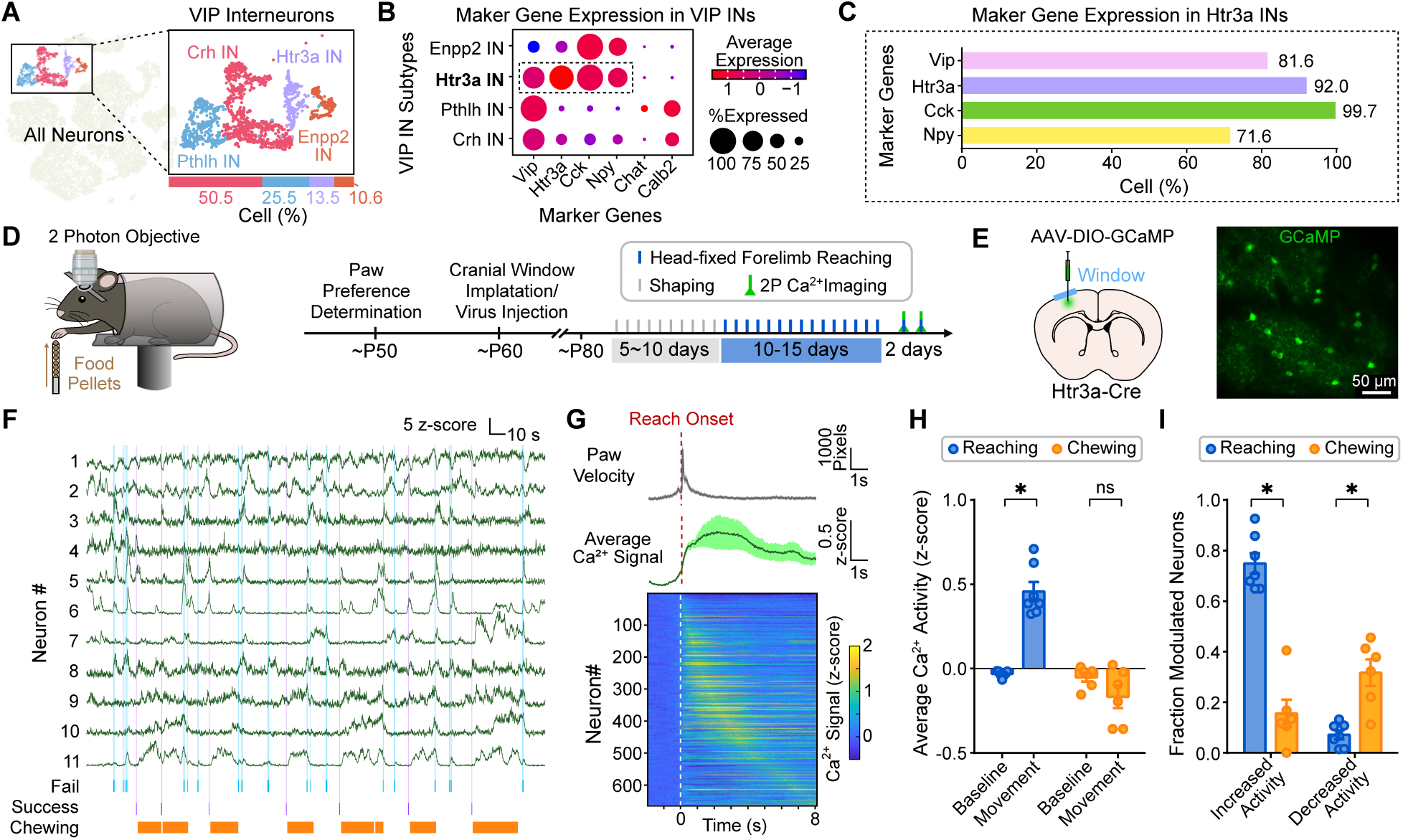
Htr3a-expressing interneurons are specifically activated during learned reaching behavior. (A) VIP-expressing interneurons in the motor cortex (highlight in color). Inset: distribution of four VIP interneuron subtypes. (B) Dot plot showing average expression levels of canonical VIP interneuron (VIP IN) markers across the four VIP subtypes. (C) Proportion of Htr3a-expressing interneurons (Htr3a IN) co-expressing *Vip*, *Htr3a, Cck* and *Npy*. (D) Schematic of the head-fixed forelimb reaching task (left) and timeline of behavior and 2-photon Ca^2+^ imaging (right). (E) Illustration of virus injection and cranial window implantation (left) and representative image showing GCaMP signal in the motor cortex (right). (F) Example fluorescence traces of 11 individual Htr3a neurons. Shaded bars indicate periods of failed reaches, successful reaches, and chewing. (G) Ca^2+^ activity of Htr3a interneurons during the reaching task. Top: Average forelimb velocity across all reaching bouts from 7 mice (mean ± SEM). Middle: Average Ca^2+^ signals of all Htr3a neurons aligned to reach onset (mean ± SEM). Bottom: Heatmap showing of Ca^2+^ activity of individual Htr3a neurons ordered by peak activation time. Dashed line indicates reach onset. (H) Quantification of average Ca^2+^ signals at baseline and during paw movement in reaching and chewing. Bars represent group means. Dots indicate individual mice. n = 7 mice for reaching; n= 6 mice for chewing. Error bars, SEM. Wilcoxon matched pairs signed rank test. *p < 0.05; ns, non-significant. (I) Proportion of Htr3a neurons modulated by reaching and chewing. Bars represent mean values; dots represent individual mice. Error bars, SEM. n = 7 mice for reaching; n= 6 mice for chewing. Wilcoxon matched pairs signed rank test. *p < 0.05; ns, non-significant. See also Figure S4 Sun et al., *BioRxiv.* 2025 | Page 15

To validate our TRAP-tdTom findings in interneurons, we further investigated the activity of Htr3a INs during motor behavior. We adapted the freely moving forelimb reaching task into a head-fixed version (Figure 3D; see method), enabling simultaneous two-photon calcium imaging of neuronal activity. Mice were trained to reach and grasp food pellets with their dominant forelimb over 10-15 days following habituation and shaping. To monitor Htr3a INs activity *in vivo*, we expressed Cre-dependent GCaMP Ca^2+^ sensors (AAV-GCaMP6s/8m or Ai148 transgenic) in Htr3a-Cre mice and implanted a cranial window over the motor cortex contralateral to their preferred paw (Figure 3E). We conducted two-photon Ca^2+^ imaging in expert mice across two daily sessions while simultaneously recording behavior using a high-speed infrared camera.

Individual Htr3a INs exhibited dynamic Ca^2+^ activities aligned with forelimb reaching (Figure 3F). Analysis of fluorescence signals aligned to reach onset revealed a robust increase in average activity following movement initiation compared to baseline (Figure 3G-3H). This increase in activity was specific to the reaching behavior, as such an increase was not observed during unrelated movements, such as chewing (Figure 3H and S4B). Approximately 75% of Htr3a INs showed significantly increased activity following reach onset, while only 7% exhibited decreased activity. In contrast, only 15% were activated during chewing, and 32% showed decreased activity following chew onset, likely resulted from the increased activity during forelimb reaches for pellets that occurred immediately preceding chewing (Figure 3I). Importantly, the activity of Htr3a INs did not differ between successful reaches and failed attempts. Both conditions elicited significant increases in post-onset activity, with similar proportions of increased or decreased activity (Figure S4C and S4D). These results suggest that Htr3a INs are selectively engaged during the execution of stereotyped, learned reaching behaviors, rather than encoding reward expectation or outcome evaluation. Taken together, these findings identify Htr3a INs as a transcriptionally distinct subtype of Cck+ VIP interneurons in the motor cortex that are preferentially activated during the performance of learned motor behaviors, implicating them in the execution phase of motor memory.

### Motor learning distinctly regulates gene expression in TRAP neurons of motor cortex and dorsal striatum

The formation and consolidation of long-term memory requires both activity-dependent transcription and *de novo* protein synthesis ^38^. To determine whether motor learning induces transcriptional remodeling in behaviorally activated neurons, we compared the transcriptomes of TRAP-tdTom neurons from early and late TRAP mice to those from control TRAP mice, within each neuronal subtype. Among ∼23,000 detected genes, differentially expressed genes (DEGs) were defined as those showing at least a 20% change in expression (|log_2_ (fold change)| > 0.26) with an adjusted p-value < 0.05 (Wilcoxon Rank Sum test followed by Holm– Bonferroni correction). To ensure robustness, we excluded genes detected in fewer than 10% of cells per subtype, genes differing across control batches, and genes differentially expressed between TRAP and non-TRAP cells in control animals – minimizing batch effects and non-learning related activity.

We first quantified DEGs across neuronal subtypes to identify cell types most affected by motor learning. In motor cortex, DEGs were predominantly enriched in six projection neuron subtypes (L2/3 IT, L4/5 IT, three L5/6 ITs, L6 CT) and one interneuron subtype (Lamp5 IN) (Figure 4A, 4B, S5A and Table S1). These neurons exhibited both upregulated and downregulated DEGs, with an overall increase in DEG numbers from early to late stages. Cross-stage comparisons showed that most DEGs were unique to a specific cell type (Figure 4C) (diagonal blocks vs other blocks), but some overlapped within a learning stage (bold-bordered blocks) across multiple subtypes, while fewer DEGs were shared between early and late stages (top right square). These findings indicate both cell type-specific and stage-dependent transcriptional remodeling in M1 during motor learning.

**Figure 4.**
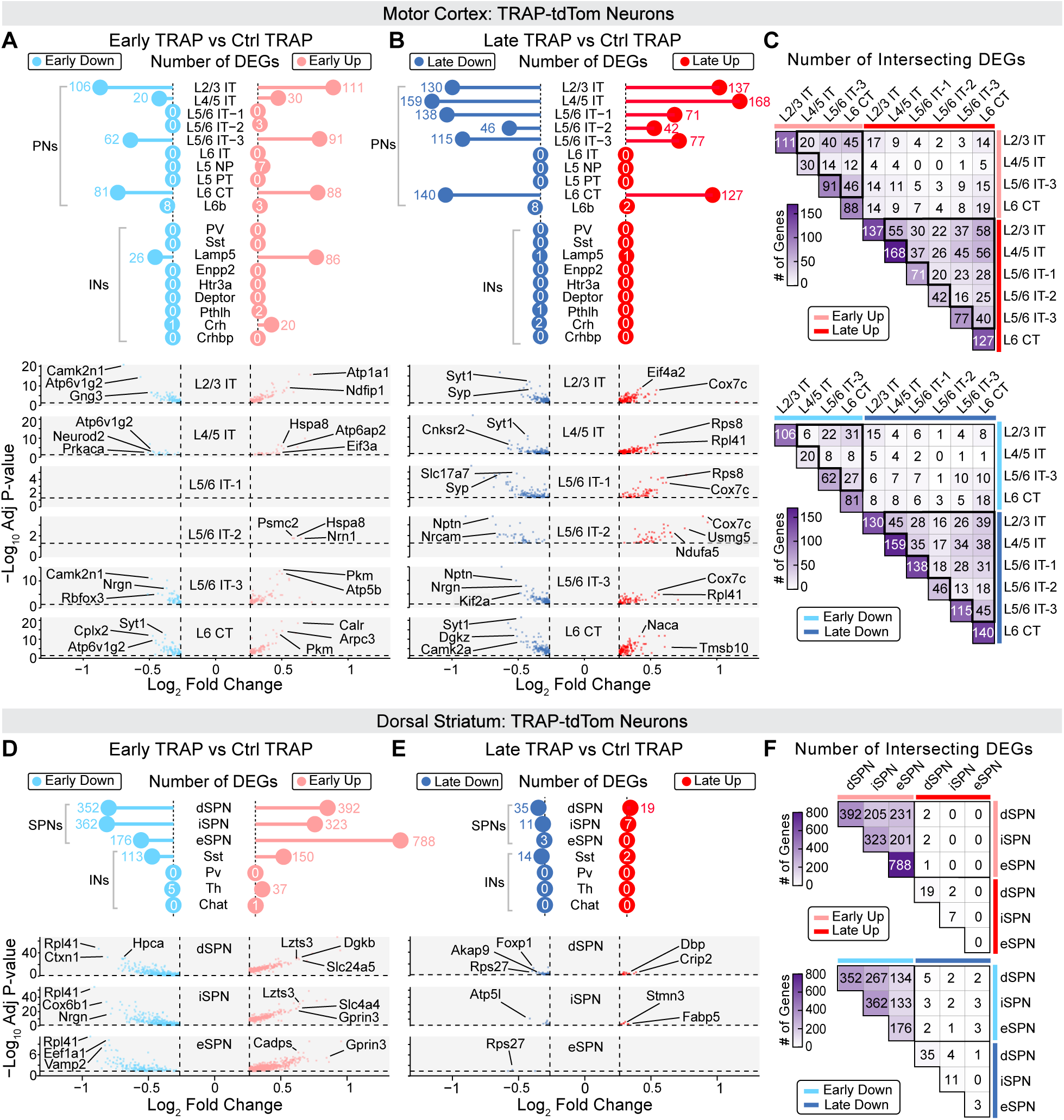
Distinct transcriptomic remodeling induced by motor learning in the motor cortex and dorsal striatum. (A) Differentially expressed genes (DEGs) identified in Early TRAP across neuronal subtypes in the motor cortex. Top, Lollipop plot displaying the number of upregulated and downregulated DEGs. Bottom, Volcano plots showing the Log_2_ fold change and -log_10_ adjusted p value of all DEGs. Wilcox rank sum test followed by Holm-Bonferroni correction. DEG Cutoff: Adjusted p-value < 0.05 and |Log_2_ Fold Change| > 0.26. (B) DEGs identified in Late TRAP across neuronal subtypes in the motor cortex. Format and statistical criteria as in (A). (C) Overlap of DEGs across neuronal subtypes in the motor cortex. Top, Heatmap showing the number of intersecting upregulated DEGs. Bottom, Heatmap showing the number of intersecting downregulated DEGs. (D) DEGs in Early TRAP across neuronal subtypes in the dorsal striatum. Top: lollipop plot of up-and downregulated DEGs. Bottom: volcano plots of log₂ fold changes and significance values. Statistical test and cutoff as in (A). (E) DEGs in Late TRAP across neuronal subtypes in the dorsal striatum. Format and criteria as in (D). (F) Overlap of DEGs between neuronal subtypes in the dorsal striatum. Top, Heatmap showing the number of intersecting upregulated DEGs. Bottom, Heatmap showing the number of intersecting downregulated DEGs. Abbreviations: IT, intratelencephalically projecting; NP, near-projecting; PT, pyramidal tract; CT, corticothalamic projecting; PV, parvalbumin; SPN, spiny projection neuron. See also Figure S5 Sun et al., *BioRxiv.* 2025 | Page 17

In contrast, DEGs in dSTR were detected predominantly in the early TRAP group (Figure 4D, 4E, S5B and Table S1). All three SPN subtypes—direct pathway SPNs (dSPNs), indirect pathway SPNs (iSPNs), and eccentric SPNs (eSPNs)—as well as Sst-expressing interneurons, showed substantial transcriptional changes. dSPN and iSPN showed ∼300-400 DEGs in both down and up regulated DEGs in early TRAP, with over 200 shared DEGs between the two major SPN types (Figure 4F), suggesting a convergent response to early learning. Eccentric SPNs, an SPN subtype that is distinguished by its unique transcriptomic profile^31^, showed the strongest transcriptional activation, with 788 upregulated and 176 downregulated DEGs. However, many eSPN DEGs overlapped with those of dSPNs and iSPNs. These findings suggest that striatal transcriptome remodeling occurs primarily during the early learning phase and is more uniform across SPN subtypes, contrasting with the delayed and subtype-divergent dynamics seen in motor cortex.

To understand the functional implications of these DEGs, we performed pathway enrichment analysis (GO and KEGG) of the identified DEGs. Shared functional pathways across M1 and dSTR included those related to synaptic plasticity, synapse assembly, protein localization, translation and energy metabolism (Figure 5A and 5B). Notably, these pathways displayed contrasting regulatory patterns between the two regions. In M1, synaptic plasticity and synapse assembly pathways were downregulated in the late TRAP neurons (late down) (Figure 5A), whereas in dSTR, these pathways were upregulated in early stage (early up) (Figure 5B). Conversely, translation and energy metabolism pathways were upregulated in M1 at both early and late training stages (Figure 5A) but were downregulated in early stage dSTR neurons (Figure 5B). Pathways related to protein localization showed mild regulation in M1 (Figure 5A) but were robustly enriched among upregulated genes in early striatal DEGs (Figure 5B). Cellular component analysis (GO) further supported these trends: in motor cortex (Figure S6A), downregulated DEGs encoded proteins localized to synaptic, dendritic, and axonal compartments, while upregulated DEGs encoded proteins localized to ribosomes and mitochondria. In contrast, the striatal neurons showed opposite pattern (Figure S5B), upregulation of genes encoding synaptic, dendritic, and axonal proteins and downregulation of ribosomal and mitochondrial components. Pathway enrichment in interneurons largely mirrored that of projection neurons, with an additional upregulation of protein processing pathways in cortical Lamp5 INs and striatal Sst INs during early learning (Figure S5A and S5B).

**Figure 5.**
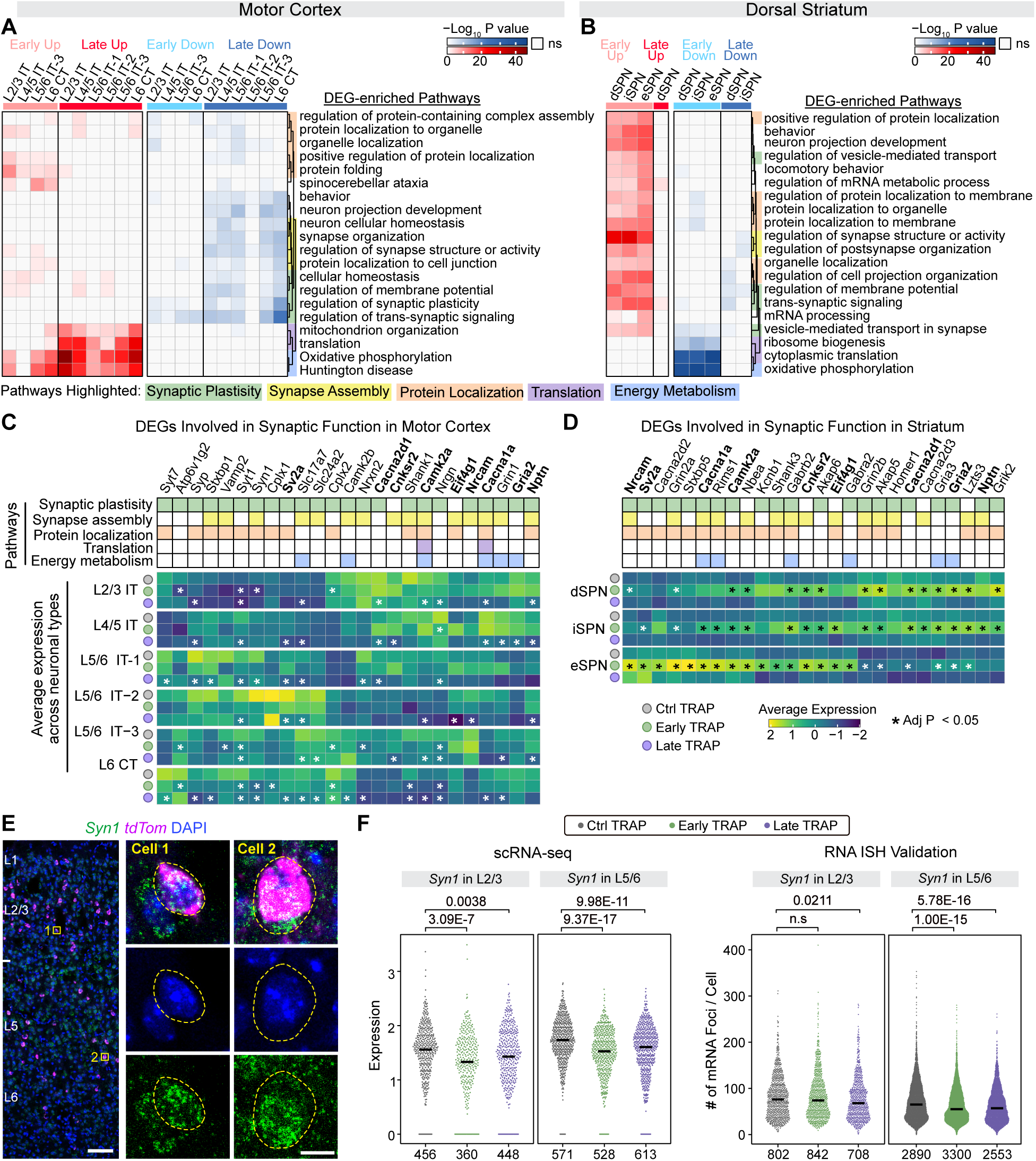
Motor learning-associated pathways are shared between motor cortex and dorsal striatum with region-and stage-specific regulation. (A) GO and KEGG pathway enrichment analysis of DEGs in projection neuron subtypes from the motor cortex. Heatmap shows -log_10_ P values, with red indicating enrichment among upregulated pathways and blue among downregulated pathways. Hypergeometric test with Benjamini-Hochberg correction. Significant terms (p < 0.01, gene count > 3, enrichment factor> 1.5) were clustered by Kappa similarity (> 0.3). Rows represent the top 20 enriched terms; columns represent neuronal subtypes. Term names are displayed in a dendrogram (right) with shaded boxes denoting their biological classifications (list at bottom). (B) GO and KEGG pathway enrichment analysis of DEGs across projection neuron subtypes in the dorsal striatum. Format, color scheme and statistics as in (A). (C) Expression of DEGs related to synaptic function in the motor cortex. Top: Table illustrating the biological categories; DEGs shared with the dorsal striatum are bolded. Bottom: Heatmaps showing average expression in Ctrl, Early, and Late TRAP neurons by subtype. Wilcox rank sum test followed by Holm-Bonferroni correction. * Adjust p value < 0.05. (D) Expression of DEGs related to synaptic function in the dorsal striatum. Top: categorization of DEGs; shared genes with the motor cortex are bolded. Bottom: heatmaps of average expression across experimental groups. Wilcox rank sum test followed by Holm-Bonferroni correction. * Adjust p value < 0.05. (E) Representative images of single-molecule RNA *in situ* hybridization (ISH) for *Syn1*in the motor cortex. Insets show enlarged views of two TRAP neurons. Yellow dashed lines outline the mask areas used for RNA puncta quantification. Scale bars: M1, 100 μm; enlarged views, 10 μm. (F) Scatter violin plots showing the expression of *Syn1* in Ctrl, Early, and Late TRAP neurons. Left: scRNA-seq data. Right: RNA ISH validations. Scatter points represent cells. Black crossbars indicate group medians. Wilcox rank sum test. P values are displayed above each violin plot; cell numbers are shown below. ns, non-significant. Abbreviations: IT, intratelencephalically projecting; NP, near-projecting; PT, pyramidal tract; CT, corticothalamic projecting; PV, parvalbumin; SPN, spiny projection neuron. See also Figure S5 and S6 Sun et al., *BioRxiv.* 2025 | Page 19

To further characterize molecular programs supporting synaptic remodeling, we focused on individual DEGs associated with synaptic function. We identified 192 downregulated DEGs in motor cortex and 264 upregulated DEGs in striatum (Figure 5C and 5D). Heatmap analysis across neuronal types confirmed region-specific expression patterns in motor cortex and striatum. These DEGs included genes encoding calcium channels (*Cacna1a*, *Cacna2d1*, *Cacna2d3* and *Cacna2d2*), and neurotransmitter receptors (*Gria2*, *Grin1*, *Gria3*, *Grin2a*, *Grin2b*, *Grik2*, *Slc17a7*, *Gabra2*, *Gabrb2*), and regulators of synaptic vesicle cycling (*Sv2a*, *Napa*, *Atp6v1g2*, *Syp*, *Stxbp1*, *Stxbp5*, *Syt1*, *Syn1*, *Cplx1*, *Cplx2*, *Vamp2* and *Rims1*). Expression changes of selected DEGs were further independently validated using single-molecule RNA *in situ* hybridization (smFISH) (Figure 5E, 5F and S6C).

Together, these results reveal distinct patterns of transcriptional remodeling in cortical and striatal neurons during motor learning. Motor cortex exhibits gradual, stage-specific molecular adaptation that persists into later learning phases, while dorsal striatum undergoes early and widespread transcriptional changes that are largely restricted to the initial acquisition phase. These region-and cell type-specific programs may underlie the temporal and functional specialization of motor learning circuits.

### Transcriptomic remodeling of glial cells is temporally synchronized with neurons during motor learning

Glial cells, including oligodendrocyte progenitor cells (OPC), oligodendrocytes (OL), astrocytes (AS) and microglia (MG) are increasingly recognized as active regulators of neural circuit plasticity during motor learning. They contribute to diverse processes such as neuronal activity modulation^16,17^, activity-dependent myelination^18,19^, and synaptic remodeling through spine formation and pruning^22,23^. These processes are likely coordinated with neuronal activity, yet the molecular basis of neuron-glia interactions during motor learning remains poorly defined.

To investigate glial contribution at the transcriptional level, we performed DEG analysis across OPCs, OLs, astrocytes, and microglia from M1 and dSTR in control, early and late learning groups (Figure S7A, S7B, S7C, S7D and Table S2). DEGs were detected in all four glial types, with patterns that mirrored those observed in neurons: in the motor cortex, DEGs were more abundant during the late learning stage, whereas in the striatum, DEGs predominated during early learning (Figure 6A and 6B; see also Figure 4A). This temporal correspondence suggests that glial transcriptional remodeling is synchronized with neuron-intrinsic gene expression changes during motor learning.

**Figure 6.**
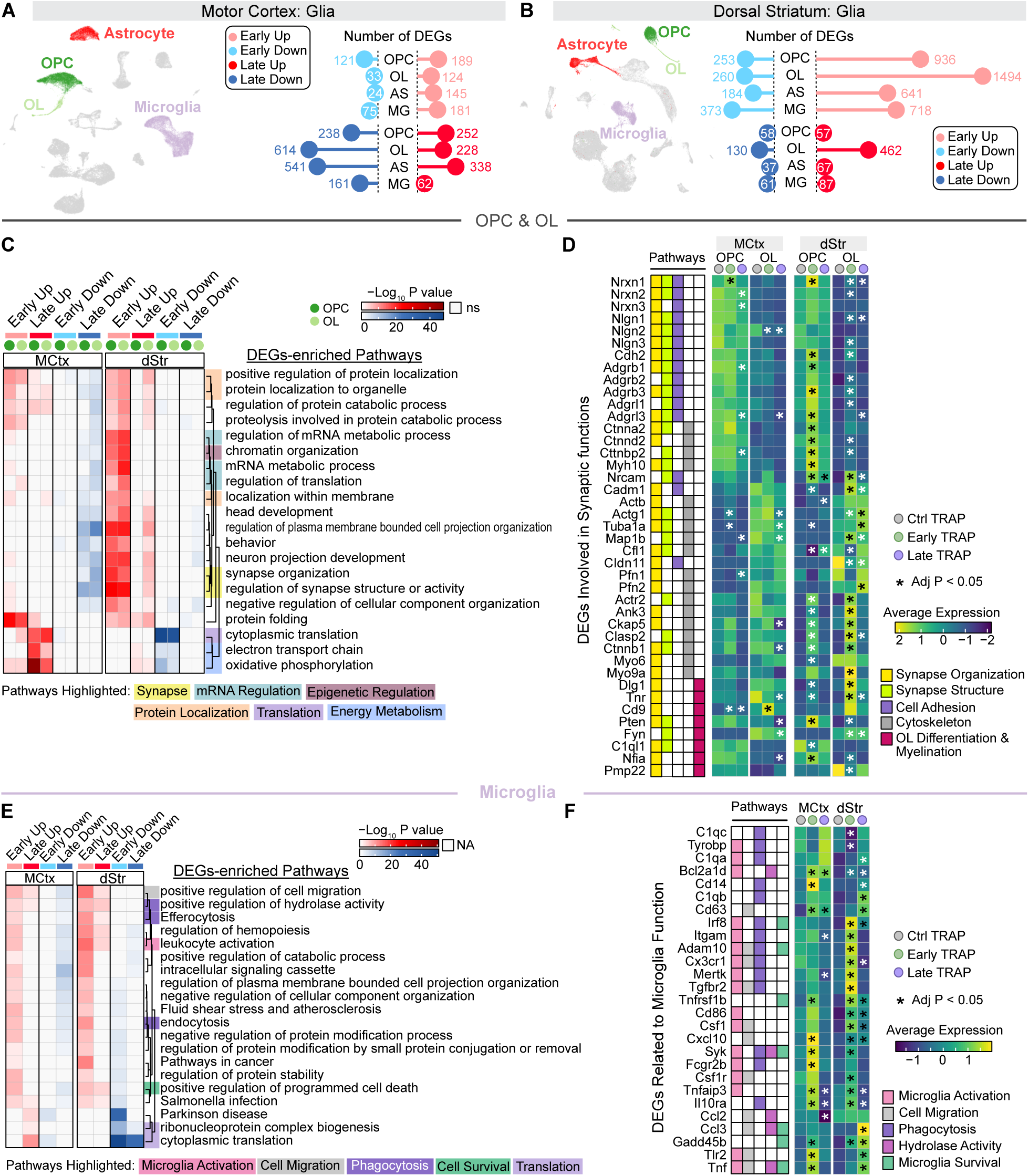
Motor learning induces transcriptomic remodeling in glial cells. (A) DEGs in glial cells of the motor cortex. Left, t-SNE plot highlights the four glia cell types. Right, Lollipop plots show the number of upregulated and downregulated DEGs in Early and Late TRAP groups compared to Ctrl. Wilcox rank sum test followed by Holm-Bonferroni correction. DEG Cutoff: Adjusted p-value < 0.05 and |Log_2_ Fold Change| > 0.26. (B) DEGs in glial cells of the dorsal striatum. Left, t-SNE plot of four glia cell types. Right: lollipop plots of DEG counts in Early and Late TRAP groups vs. Ctrl. Statistical method and cutoff as in (A). (C) GO and KEGG pathway enrichment analysis of DEGs in OPCs and OLs. Heatmap displaying -log_10_ P values with red indicating upregulated pathways and blue indicating downregulated pathways. Hypergeometric test with Benjamini-Hochberg correction. Significant terms (p < 0.01, gene count > 3, enrichment factor > 1.5) were clustered by Kappa similarity (> 0.3), and the top 20 enriched pathways are shown. Term names are displayed in a dendrogram (right) with shaded boxes denoting their biological classifications (list at bottom). (D) Expression of DEGs associated with synaptic functions in OPCs and OLs. Left: Table illustrating the biological categories of each DEGs. Right: Heatmaps showing average expression across Ctrl, Early, and Late TRAP neurons in OPCs and OLs of the motor cortex and dorsal striatum. Wilcox rank sum test followed by Holm-Bonferroni correction. * Adjust p value < 0.05. (E) GO and KEGG pathway enrichment analysis of DEGs in microglia. Heatmap formatting, organization and statistics as in (C). (F) Expression of DEGs related to microglial functions. Top: Table illustrating the biological categories of each DEGs. Bottom: Heatmaps showing average expression in Ctrl, Early, and Late TRAP groups in microglia from the motor cortex and dorsal striatum. Wilcox rank sum test followed by Holm-Bonferroni correction. * Adjust p value < 0.05. Abbreviations: OPC, oligodendrocyte precursor cell; OL, oligodendrocyte; AS, astrocyte; MG, microglia; MCtx, motor cortex; dSTR, dorsal striatum. See also Figure S7

Shared pathways across neurons and glia also included translation, energy metabolism, and protein localization (Figure 6C, 6E and S7E), suggesting that some transcriptional programs are regulated in a stage-dependent manner regardless of cell identity. However, glia also displayed cell type-specific molecular signatures. In OPCs and OLs (Figure 6C and 6D), pathways related to synapse organization and structure were also enriched, as in neurons, but the specific genes contributing to these pathways were different. These DEGs included genes involved in cell-cell adhesion (e.g., *Neurexins* and *Neuroligins*), cytoskeletal components (e.g., *Actin*, *Catenins*, and *Tubulin*), and oligodendrocyte differentiation (e.g., *Dlg1*, *Tnr*, *Cd9*, *Pten*, and *Fyn*), all of which may contribute to activity-dependent myelination. OPCs and OLs, as well as astrocytes (Figure 6C and S7E), also showed significant enrichment of DEGs in pathways related to mRNA processing and epigenetic regulation, including mRNA splicing, transport, stability, microRNA regulation, and chromatin organization (Figure S7F). These changes were more prominent in motor cortex during late learning and in striatum during early learning, consistent with region-and stage-specific dynamics.

In microglia, DEGs were enriched in pathways associated with canonical microglia functions, such as microglia activation, migration, phagocytosis and survival (Figure 6E and 6F). Notably, these genes were upregulated ruing early learning stage, coinciding with the peak dendritic spine formation in the motor cortex ^5–9^. These findings are consistent with prior evidence that microglia are required for learning-induced synaptogenesis and that their depletion impairs motor learning^22^. Together, these results underscore the importance of oligodendrocytes and microglia in circuit remodeling and further suggest that distinct glial populations contribute to learning through temporally coordinated, transcriptionally regulated programs.

## Discussion

Motor learning induces activity-dependent structural and molecular adaptations in the primary motor cortex and dorsal striatum, but the underlying transcriptional mechanisms remain incompletely understood. In this study, we combined activity-dependent genetic labeling (TRAP) strategy with single-cell RNA sequencing (scRNA-seq) to define the transcriptomic landscape of behaviorally engaged neuronal and glial populations during a forelimb reaching task (Figure 1). Our results reveal that diverse neuronal subtypes, particularly interneurons, are dynamically recruited throughout learning (Figure 2). Notably, we identified Htr3a-expressing interneurons (Htr3a INs) in the motor cortex, a previously underappreciated subset of VIP interneurons, as specifically activated during reaching. Two-photon Ca^2+^ imaging confirmed their activation was tightly time-locked to movement execution, but not outcome evaluation, suggesting a role in fine-tuning cortical output *via* local disinhibition (Figure 3). Transcriptomic profiling further revealed distinct, region-and stage-specific molecular programs across projection neurons and glia, implicating pathways related to synaptic remodeling, translation, energy metabolism, and protein localization (Figure 4 and 5). Furthermore, significant transcriptomic remodeling was detected in glial cells, which exhibited stage- and region-specific patterns similar to neuronal ensembles (Figure 6).

A major finding is the identification of Htr3a INs as a transcriptionally and functionally distinct subtype of VIP interneurons that are selectively engaged during the execution of learned motor behavior. Unlike previously described VIP interneurons, which exhibit preparatory activity and are thought to facilitate disinhibition during motor planning ^14^, Htr3a INs were recruited after reach onset. These cells co-express Cck, a marker of interneurons capable of directly targeting excitatory pyramidal neurons, suggesting they may exert more direct control over output neurons compared to canonical VIP INs that preferentially inhibit other interneurons^39,40^. This raises key questions regarding their downstream targets and circuit integration: Do Htr3a INs modulate projection neurons directly, or do they shape behavior through interactions with SST or PV interneurons? Future studies using cell-type–specific manipulations and functional connectomics will be critical to define their role in motor cortical microcircuits. Notably, Htr3a is a ligand-gated ionotropic serotonin receptor. Its activation by 5-hydroxytryptamine (5-HT) is known to directly depolarize neurons, implying that serotonergic signaling may potently excite Htr3a INs during learning. This raises the intriguing possibility that 5-HT plays a direct, modulatory role in enhancing Htr3a IN recruitment and motor circuit plasticity. Future studies exploring serotonergic input to these neurons may uncover a powerful neuromodulatory axis linking behavioral state, interneuron excitability, and learning.

We also found that transcriptomic remodeling in projection neurons varies markedly between M1 and dSTR. In M1, synaptic genes were persistently downregulated from early to late learning, consistent with reports of learning-induced spine elimination and pruning in pyramidal neurons^5–8^. In contrast, SPNs in the striatum showed robust upregulation of synaptic genes during early learning, with little change at later stages. This pattern suggests that striatal neurons undergo an early, transient transcriptional activation, whereas cortical neurons exhibit slower, more sustained reorganization. While the striatum’s involvement in motor learning has been established at the circuit level ^4^, our data highlight a previously underappreciated temporal asymmetry in molecular plasticity across these regions. These differences may reflect the unique computational roles of cortex and striatum during motor skill acquisition, such as top-down command encoding versus outcome gating and habit formation.

In parallel, glial populations also displayed substantial transcriptional changes that were tightly synchronized with neuronal adaptations. Oligodendrocytes and OPCs exhibited dynamic regulation of genes involved in cell-cell adhesion, cytoskeletal remodeling and myelination, pointing to their potential roles in neuronal plasticity through activity-dependent myelination. Microglia showed early and transient activation of genes linked to migration, phagocytosis, and survival, consistent with their involvement in learning-induced synaptic plasticity^22,23^. Furthermore, astrocytes, OPCs, and oligodendrocytes showed significant enrichment in pathways related to mRNA processing and chromatin remodeling, suggesting active post-transcriptional and epigenetic reprogramming during learning. Transcriptomic reprogramming in glia has also been reported in non-motor behavioral contexts^41,42^, highlighting the broader engagement of glia in plasticity beyond motor learning. An outstanding question is how glia and neurons communicate at the molecular level during learning. Our findings set the stage for spatially resolved approaches, such as spatial transcriptomics combined with TRAP labeling, to map interactions between activated neurons and nearby glia. This strategy could reveal molecular signaling mechanisms that coordinate plasticity across cell types and could be expanded to other behaviors or disease models to assess whether these programs are conserved or perturbed.

In summary, our study provides a comprehensive molecular atlas of the learning-activated motor circuit, uncovering cell-type–specific transcriptional programs across excitatory neurons, interneurons, and glial populations. These findings underscore the complexity of motor memory formation and reveal the layered contributions of cortical and striatal populations to learning-related plasticity. Moving forward, integrating transcriptomic, structural, and functional approaches will be essential for dissecting how these cellular programs converge to support adaptive behavior—and may ultimately inform interventions for motor dysfunction in neurological disorders.

## Resource availability

### Lead contact

Further information and requests for resources and reagents should be directed to and will be fulfilled by the lead contact, Jun B. Ding.

### Materials availability

This study did not generate new unique reagents.

### Data and code availability

Single cell RNA sequencing data have been deposited to Gene Expression Omnibus (GSE300408) and are publicly available as of the date of publication. Any additional information required to reanalyze the data reported in this paper is available from the lead contact upon request.

## Acknowledgments

We thank Dr. Ivan Soltesz for generously sharing the Htr3a-Cre line. We thank Dr. Yu-Wei Wu for his valuable input in conception of this study, Dr. Dongli Xu and Dr. Xiaobai Ren for technical support, and all members of Ding lab for valuable discussions. Sequencing was conducted by the Genome Sequencing Service Center (GSSC) at Stanford University. This study was supported by grants from the Stanford School of Medicine Dean’s Postdoctoral Fellowship (to Y.S and R.H.R), Brain & Behavior Research Foundation Young Investigator Grant (to Y.S), NINDS/NIH K99NS130078 (to R.H.R.), Parkinson’s disease foundation postdoc fellowship PF-PRF-1264715 (to F.J.H), NINDS/NIH R01NS091144 (to J.B.D.), GG gift fund (to J.B.D.), Deng family gift fund (to J.B.D.), Catalyst grant from The Phil & Penny Knight Initiative for Brain Resilience at the Wu Tsai Neurosciences Institute, Stanford University (to J.B.D).

## Author contributions

Y.S., S.W., and J.B.D. conceived the project and designed experiments. Y.S. performed the scRNA-seq and single-molecule RNA ISH experiments and analyzed data under S.W.’s guidance. Y.S. and F.J.H. conducted immunostaining and cell counting. R.H.R. and Y.S. performed head-fixed reaching and calcium imaging and analyzed data. Y.S. and J.B.D. wrote the manuscript with input from all authors.

## Materials and methods

Key resources table

Key resources table
| REAGENT or RESOURCE | SOURCE | IDENTIFIER |
| --- | --- | --- |
| **Antibodies** |  |  |
| Anti-c-Fos antibody | Synaptic Systems | Cat# 226 003; RRID:AB_2231974 |
| Donkey anti-Rabbit IgG (H+L) Secondary Antibody, Alexa Fluor 488 | Invitrogen | Cat# A-21206 |
| **Bacterial and virus strains** |  |  |
| AAV1-syn-FLEX-jGCaMP8m-WPRE | GENIE Project | Addgene viral prep # 162378-AAV1; RRID:Addgene_162378 |
| AAV5.CAG.Flex.GCaMP6s.WPRE.SV40 | GENIE Project | Addgene viral prep # 100842-AAV5; RRID:Addgene_100842 |
| AAV1-syn-FLEX-jGCaMP8m-WPRE | GENIE Project | Addgene viral prep # 162378-AAV1; RRID:Addgene_162378 |
| **Chemicals, peptides, and recombinant proteins** |  |  |
| 4-Hydroxytamoxifen (4-OHT) | Sigma-Aldrich | Cat# H6278 |
| Kynurenic acid | Sigma-Aldrich | Cat# K3375 |
| Nω-Nitro-L-arginine | Sigma-Aldrich | Cat# N5501 |
| L-Glutathione reduced | Sigma-Aldrich | Cat# G4251 |
| Sodium pyruvate | Sigma-Aldrich | Cat# P2256 |
| Papain | Worthington | Cat# LK003176 |
| **Critical commercial assays** |  |  |
| Dead Cell Removal Kit | Miltenyi Biotec | Cat# 30-090-101 |
| Chromium Single Cell 3' Library & Gel Bead Kit v2, 4 rxns | 10x Genomics | Cat# PN-120267 |
| Chromium Single Cell 3' Library & Gel Bead Kit v3, 4 rxns | 10x Genomics | Cat# PN-1000092 |
| RNAscope Multiplex Fluorescent Detection Kit v2 | Advanced Cell Diagnostics (ACD) | Cat# 323110 |
| RNAscope probe: tdTomato | ACD | Cat# 317041 |
| RNAscope probe: tdTomato-C2 | ACD | Cat# 317041-C2 |
| RNAscope probe: tdTomato-C3 | ACD | Cat# 317041-C3 |
| Mm-Fos-C2 | ACD | Cat# 316921-C2 |
| Mm-Arc-C3 | ACD | Cat# 316911-C3 |
| Mm-Syt7 | ACD | Cat# 546201 |
| Mm-Vamp2-C3 | ACD | Cat# 573151-C3 |
| Mm-Syn1 | ACD | Cat# 445241 |
| Mm-Snca-C2 | ACD | Cat# 313281-C2 |
| mm-Park7 | ACD | Cat# 514201 |
| TSA Plus Fluorescein | Akoya Biosciences | Cat# NEL741001KT |
| TSA Plus Cyanine 3 | Akoya Biosciences | Cat# NEL744001KT |
| TSA Plus Cyanine 5 | Akoya Biosciences | Cat# NEL745001KT |
| **Deposited data** |  |  |
| Single cell RNA-seq dataset | This paper | GSE300408 |
| **Experimental models: Organisms/strains** |  |  |
| Mouse: B6.129(Cg)-Fos<sup>tm1.1(cre/ERT2)Luo/J</sup> (Fos-Cre<sup>ER</sup>) | The Jackson Laboratory | JAX# 021882; RRID:IMSR_JAX:021882 |
| B6.129(Cg)-Arc<sup>tm1.1(cre/ERT2)Luo/J</sup> (Arc-Cre<sup>ER</sup>) | The Jackson Laboratory | JAX# 021881; RRID:IMSR_JAX:021881 |
| Mouse: B6.Cg-Gt(ROSA)<sup>26Sortm9</sup>(CAG-tdTomato)Hze/J (Ai9) | The Jackson Laboratory | JAX# 007909; RRID:IMSR_JAX:007909 |
| Mouse: B6.Cg-Gt(ROSA)<sup>26Sortm14</sup>(CAG-tdTomato)Hze/J (Ai14) | The Jackson Laboratory | JAX# 007914; RRID:IMSR_JAX:007914 |
| Mouse: B6.Cg-Igs7<sup>tm148.1</sup>(tetO-GCaMP6f,CAG-tTA2)Hze/J (Ai148) | The Jackson Laboratory | JAX# 030328; RRID:IMSR_JAX:030328 |
| Mouse: B6.FVB(Cg)-Tg(Htr3a-cre)NO152Gsat/Mmucd (Htr3a-Cre) | MMRRC | 037089-UCD; RRID:MMRRC_037089-UCD |
| Mouse: Fos-Cre<sup>ER::</sup> Ai9 (Fos-TRAP) | This paper | N/A |
| Mouse: Fos-Cre<sup>ER::</sup> Ai14 (Fos-TRAP) | This paper | N/A |
| Mouse: Arc-Cre<sup>ER::</sup> Ai9 (Arc-TRAP) | This paper | N/A |
| Mouse: Arc-Cre<sup>ER::</sup> Ai14 (Arc-TRAP) | This paper | N/A |
| Mouse: Htr3a-Cre:: Ai148 | This paper | N/A |
| Software and algorithms |  |  |
| ImageJ | Schneider et al., 2012 | https://imagej.net; RRID:SCR_003070 |
| CellProfiler |  | https://cellprofiler.org; RRID:SCR_007358 |
| MATLAB | Mathworks | https://www.mathworks.com; RRID:SCR_001622 |
| Prism 9 | GraphPad Software | http://www.graphpad.com; RRID:SCR_002798 |
| Affinity Designer | Serif | https://affinity.serif.com/en-us/designer/; RRID:SCR_016952 |
| R (v4.2.3) | N/A | http://www.r-project.org; RRID:SCR_001905 |
| Cell Ranger (v3.1.0) | 10x Genomics | https://support.10xgenomics.com/single-cell-gene-expression/software/pipelines/latest/what-is-cell-ranger; RRID:SCR_017344 |
| Seurat (v4) | Hao et al., 2021 | https://satijalab.org/seurat/get_started.html; RRID:SCR_016341 |
| DropViz | Saunders et al., 2018 | http://dropviz.org/; RRID: N/A |
| Cell Type Knowledge Explorer | BRAIN Initiative Cell Census Network (BICCN) | https://knowledge.brain-map.org/celltypes; RRID:SCR_022793 |
| Metascape | Zhou et al., 2019 | http://metascape.org/gp/index.html#/main/step1; RRID:SCR_016620 |
| EnhancedVolcano | Blighe K et al., 2024 | https://bioconductor.org/packages/EnhancedVolcano/; RRID:SCR_018931 |
| pheatmap | Kolde R., 2018 | https://www.rdocumentation.org/packages/pheatmap/versions/0.2/topics/pheatmap; RRID:SCR_016418 |
| DeepLabCut (v2.1.0) | Mathis et al., 2018 | https://github.com/DeepLabCut/DeepLabCut; RRID:SCR_021391 |
| BORIS (v7.13.4) | Friard O and Gamba M., 2016 | https://www.boris.unito.it/; RRID:SCR_025700 |
| Custom scripts for analysis | This paper |  |
| Other |  |  |
| Food pellets (10 mg) | TestDiet | Cat# 1811328 |
| Food pellets (20 mg) | Bio-Serv | Cat# F0071 |

## Experimental model and subject details

### Animals

All procedures were conducted in accordance with NIH guidelines and were approved by the Institutional Animal Care and Use Committee (IACUC) at Stanford University. The mouse lines used in this study were bred in-house on a C57BL/6 background and are listed in the key resource table. Both male and female mice (> 6 weeks of age) were used for all experiments, with littermates randomly assigned to experimental groups. Mice were group-housed under a standard 12-hour light-dark cycle with *ad libitum* access to food and water, except during behavioral training. One week before training, mice were switched to a reverse 12-hour light-dark cycle, allowing training in their active (dark) phase. All training sessions were conducted at consistent times each day to minimize circadian variability. For food pellet reaching task, mice were food restricted to ensure motivation. Mice were weighed to obtain baseline bodyweight and then restricted to ∼0.1 g food/1 g body weight per day. After 2 days of food restriction, their body weight reduced to 85%∼90% of the baseline weight and was maintained throughout the training.

## Method details

### Free moving forelimb reaching task

Mice were trained to reach single food pellet using their dominate limbs through a narrow opening in front of a behavior chamber (20 cm height, 15 cm depth, 8.5 cm width with a 0.5 cm slit opening) under food restriction, as previously described ^8,9^. The training process includes 3∼5 days of shaping followed by 8 days of training. During the shaping phase, mice were gradually habituated to the training environment, and their dominant limb was identified. On the first day, two mice were placed in the chamber for 20 minutes with 40 food pellets (TestDiet, Cat# 1811328) provided on the floor and the next day, mice were placed individually with 20 food pellets. On the third day, to identify their dominant limb, a tray filled with pellets was positioned outside the chamber wall, requiring mice to reach through the opening to obtain food pellets. Each mouse was placed in the chamber individually for 20 minutes or until it attempted 20 reaches. Shaping was considered complete when a mouse made either five consecutive reaching attempts with one limb or at least 70% of its total attempts with the same limb. Mice failing to meet this criterion after 5 days were excluded from the experiment. The training phase lasted eight consecutive days, during which shaped mice were individually placed in the chamber and trained to reach and grasp single food pellets from a platform using their dominant paw. Each session lasted for 30 reaches or 20 minutes, whichever came first, and performance was monitored. Reaching outcomes were classified as success (grasped, retrieved and ingested pellet), fail (missed, displaced or dropped pellet), in-vain reach (attempts when no pellet was available), and contralateral reach (attempts with the non-dominant limb). The success rate was calculated as the number of successful reaches divided by the sum of successful and failed reaches, excluding in-vain and contralateral attempts. Early TRAP mice were only trained for two consecutive days and late TRAP mice were trained for eight consecutive days. Ctrl TRAP mice followed the same training timeline as early or late TRAP groups but were only exposed to the training chamber without training for reaching and were given 30 food pellets in the chamber. For memory retrieval experiments, mice underwent an additional training session (or chamber exposure for Ctrl) three days after their final training day.

### Tamoxifen preparation and administration

4-Hydroxytamoxifen (4-OHT) solution was prepared by dissolving 4-OHT powder (Sigma, Cat#H6278) in ethanol at 100 mg/ml and sonicating at 55 °C for 15 min, then aliquoted and stored at -20 °C until use. Before injection, 4-OHT aliquots were reheated to 55 °C with sonication until clear, then diluted with sunflower seed oil to a final concentration of 10 mg/ml. To induce Cre-dependent tdTomato expression in activated neurons, 4-OHT was injected intraperitoneally at 50 mg/kg body weight following training. Early TRAP mice received 4-OHT on training day 1 and 2, and late TRAP mice received it on training day 7 and 8. Ctrl TRAP mice were given 4-OHT following the same schedule as either early or late TRAP mice.

### Single-cell isolation, cDNA library construction and sequencing

Mice were anaesthetized with isoflurane and perfused with cold artificial cerebrospinal fluid (ACSF) containing 125 mM NaCl, 2.5 mM KCl, 1.25 mM NaH_2_PO_4_, 25 mM NaHCO_3_, 15 mM glucose, 2 mM CaCl_2_ and 1 mM MgCl_2_, oxygenated with 95% O_2_ and 5% CO_2_ (300∼305 mOsm, pH 7.4). Coronal sections were prepared using a vibratome (Leica, VT1200) at 300 µm thickness and subsequently incubated in ACSF for 30 min at 34 °C. Following recovery, motor cortex of Fos-TRAP mice and dorsal striatum of Arc-TRAP mice were dissected under a stereo microscope and quickly transferred to chambers containing Hanks’ Balanced salts solution with supplements (HBSS/s). The HBSS/s solution consisted of 5.3 Mm KCl, 0.4 mM KH_2_PO_4_, 138 mM NaCl, 0.34 mM Na_2_HPO_4_, 5.6 mM D-Glucose, 10 mM HEPES, 2 mM MgCl_2_, 1 mM Kynurenic acid (Sigma, Cat#K3375), 0.1 mM Nω-Nitro-L-arginine (Sigma, Cat#N5501), 5 µM L-Glutathione reduced (Sigma, Cat#G4251), 1 mM Sodium pyruvate (Sigma, Cat#P2256), and was continuously oxygenated with pure O_2_ (∼295 mOsm, pH 7.4). Tissue from either the contralateral or ipsilateral hemisphere relative to the trained limbs were pooled from two early TRAP or late TRAP mice. Tissues from two Ctrl TRAP mice are pooled by right or left hemisphere. Pooled tissue was digested with papain (20 units/ml in HBSS/s) (Worthington, Cat#LK003176) at 37 °C for 1 h with gentle stirring and continuous oxygenating. The digested tissue was then mechanically triturated using fire-polished glass capillaries and filtered through 40 um strainers. Dead cells and debris were removed using a Dead Cell Removal Kit (Miltenyi Biotec, Cat#130-090-101) following manufacturer’s instruction. Single cells were resuspended in cold PBS to a final concentration of 700∼1200 cells/ml.

cDNA libraries were prepared by Genome Sequencing Service Center (GSSC) at Stanford University using Chromium Next GEM Single Cell 3’ Library Construction Kit v2 (10x Genomics, PN-120267) or v3 (PN-1000092), following manufacture’s instruction. Briefly, single-cell suspensions were loaded into the Chromium Chip for target recovery of 10,000 cells. By running the Chip in a Chromium controller, cells were captured and lysed within droplets containing barcoding beads and incubated at 53 °C for 45 min to generate barcoded cDNAs from the poly A-tailed mRNA transcripts. Single-stranded cDNAs were then released from droplets, PCR amplified (11 cycles), fragmented, size-selected, ligated with adapters, and amplified by index PCR (11∼12 cycles) to construct the final cDNA libraries. Finally, libraries were sequenced on an Illumina HiSeq4000 (paired-end with read lengths of 100 nt).

### Immunohistology

Mouse brain tissue was collected 30–60 minutes after behavioral training for immunostaining. Mice were transcardially perfused with phosphate-buffered saline (PBS), followed by 4% paraformaldehyde (PFA) in PBS. Brains were then dissected, post-fixed in 4% PFA for 2 hours at room temperature and transferred to 30% sucrose in PBS at 4 °C until fully dehydrated (∼48 hours). The brains were subsequently embedded in OCT compound, frozen, and sectioned at 30 or 40 μm thickness using a microtome (Leica) and stored in PBS. For immunostaining, sections were rinsed three times in PBS, permeabilized with 0.5% Triton X-100 in PBS for 30 minutes, then blocked for 1 hour at room temperature in a solution containing 1% BSA, 10% normal donkey serum (NDS) and 0.1% Triton X-100 in PBS. Sections were incubated overnight at 4 °C with primary antibody (Rabbit anti-c-Fos, Synaptic Systems, Cat#226-003; 1:1000 in blocking solution), washed in PBS, and incubated with secondary antibodies (Donkey anti-rabbit Alexa 488; 1:500 in PBS with 1 % BSA) for 2 hours at room temperature. After final washes, sections were mounted on glass slides using DAPI-containing mounting medium (Vector Laboratories, Cat# H-1500). Stained sections were stored at 4 °C and imaged using a 10x /0.45 NA objective on a confocal microscope (Zeiss, LSM900).

### *In situ* RNA hybridization

Multiplex RNA *in situ* hybridization was performed on fixed-frozen mouse brain sections using the RNAscope® Multiplex Fluorescent v2 Assay (Advanced Cell Diagnostics, Cat# 323270), following the manufacturer’s protocol (Doc# 323100-USM) with minor modifications. Mice were perfused with 4% PFA in PBS, and brains were collected and post-fixed overnight at 4 °C followed by dehydrated with 30% sucrose in PBS. The tissue was frozen in OCT and sectioned at 20 μm thickness onto Superfrost Plus slides (Fisherbrand, Cat#12-550-15). Slides were baked at 60°C for 30 min, fixed in 4% PFA for 15 min at 4°C, and dehydrated through a graded ethanol series (15%, 75% and 100%, 5 mins each). Tissue sections were then treated with hydrogen peroxide (10 mins at RT), target retrieval reagent (3 mins in steam), and Protease III (20 mins at 40°C). Probes targeting RNA of interest were hybridized for 2 hours at 40 °C, followed by signal amplification (AMP1–3) and sequential TSA-based fluorophore development (Fluorescein, Cy3, and Cy5). Each channel was developed individually using HRP-conjugated probes and corresponding TSA fluorophores, with HRP blocking between rounds to prevent cross-reactivity. Sections were stained with DAPI (1:10000; Invitrogen, D3571), mounted in anti-fade medium (Southern Biotech, Cat#0100-01), and imaged using Zeiss LSM900 confocal microscopy equipped with a 20x/0.8 NA objective. A list of RNA probes used is provided in the Key Resources Table.

### Stereotaxic viral injection

Cre-dependent GCaMP6s (AAV5.CAG.Flex.GCaMP6s.WPRE.SV40; Addgene#100842-AAV5, titer ≥ 1×10^13^ vg/mL) or GCaMP8m viruses (AAV1-syn-FLEX-jGCaMP8m-WPRE; Addgene#162378-AAV1, titer ≥ 1×10^13^ vg/mL) were stereotaxically injected into the motor cortex of adult Htr3a-Cre mice (MMRRC, 037089-UCD; a gift from Dr. Ivan Soltesz, Stanford University). Mice were anesthetized with isoflurane (1.5–2.0%, 1 L/min O₂) and secured in a stereotaxic frame. A small craniotomy was made over the contralateral primary motor cortex (to the trained limb; coordinates: AP 1.0 mm, ML 1.5 mm from bregma, and DV 1 mm from brain surface). A total volume of 350-400 nl virus was injected at 50-100 nL/min using a pulled glass capillary needle (Drummond Scientific, Cat# 5-000-2005; Sutter micropipette puller). After injection, the incision was closed with 3–5 sutures. Animals were allowed to recover from surgery at least 2 weeks before behavior experiments.

### Cranial window implantation

A cranial window (3 × 3 mm, #1 coverslip) was implanted over the primary motor cortex contralateral to the trained limb to enable optical access to neuronal activity. Mice were anesthetized with isoflurane (1.5–2.0% in 1 L/min O₂), and a 3×3 mm craniotomy was made using #11 scalpel blades (Fine Science Tools). The craniotomy was centered ∼2 mm lateral to bregma. A coverslip was placed directly onto the dura and sealed to the skull using tissue adhesive (Vetbond tissue adhesive, 3M). A custom titanium headplate was then positioned over the implant and attached to the skull with dental acrylic (Metabond, Parkell). Animals were allowed to recover from surgery at least 2 weeks before behavior experiments.

### Head-fixed reaching task

Mice were trained to perform a forelimb reaching task under head fixation to enable simultaneous two-photon imaging. Food-restricted mice first underwent a habituation period lasting 3–7 days. During habituation, animals were placed in a body restraint tube that allowed full forelimb extension, while their headplates were secured to overhead bars. Food pellets (Bio-Serv, Cat# F0071) were delivered by hand using tweezers. Head-fixation duration was gradually increased based on each animal’s comfort, aiming for at least 20 minutes of calm fixation during which mice actively consumed food pellets. After habituation, mice were trained daily for a minimum of 8 consecutive days. Each daily training session (15–30 minutes) consisted of repeated trials in which single food pellets were delivered *via* a custom pellet delivery device, allowing mice to self-initiate reaches with their dominant forelimb. Initially, the pellet was positioned near the mouth to encourage licking. Once mice consistently performed three autonomous licks, the pellet holder was lowered near the dominant paw to train tactile location and grasping behavior. As mice learned to touch and grasp the pellet, the holder was moved forward (1-2 cm below the nose) to encourage full reaching movements. Reach outcomes were recorded using a high-speed infrared camera (FLIR) and classified as follows: success (pellet grasped and consumed), drop (pellet grasped but dropped before consumption), fail (missed or displaced pellet), and in-vain (reach made in the absence of a pellet; excluded from analysis).

### Two-photon calcium imaging

*In vivo* two-photon imaging was performed in Htr3a-Cre mice expressing Cre-dependent calcium indicators, using a combination of viral and transgenic strategies. Three mice were injected with AAVs encoding GCaMP6s, one with GCaMP8m, and three were bred with the Ai148 (GCaMP6f) reporter line. All mice were pre-trained on the head-fixed reaching task and imaged while actively performing the behavior. Imaging was conducted using a custom-built two-photon microscope equipped with a resonant scanner (LotosScan, Suzhou Institute of Biomedical Engineering and Technology) and a 25×/1.0 NA water-immersion objective (Olympus). GCaMP indicators were excited at 925 nm using a mode-locked, tunable ultrafast laser (InSightX3, Spectra-Physics) with laser power ranging from 15 to 80 mW at the objective. Images were acquired at 20 Hz (600 × 600 pixels) from consistent fields of view located 50 to 250 µm below the pial surface, with each session lasting for 5 minutes.

## Quantification and statistical analysis

### Immunostaining and RNA ISH quantifications

In Fos-TRAP and Arc-TRAP mice, neurons activated during learning were labeled with tdTomato (TRAP-tdTom), while neurons activated during memory retrieval were identified using anti-c-Fos antibody staining and Arc RNA ISH, respectively. Confocal images from these animals were quantified for TRAP-tdTom and c-Fos or Arc populations, as well as their overlap (“Double positive”). All analysis was performed blinded to experimental conditions to prevent bias.

### Fos-TRAP and c-Fos staining

For Fos-TRAP, two coronal brain sections (40 µm thick), located approximately 600 µm and 900 µm anterior to bregma, were selected for analysis of the primary motor cortex (M1). ROIs covering M1 was manually defined in both hemispheres in ImageJ ^43^. The “Despeckle” and “Watershed” functions were applied on tdTomato and c-Fos fluorescence channels to reduce noise and segment merged cells. Thresholds were then manually set for tdTomato and c-Fos channel, and cells were automatically identified and counted using the “Analyze Particles” function. Double positive cells were defined as tdTomato-positive cells in which more than 40% of their pixels also showed c-Fos signal.

### Arc-TRAP and Arc RNA ISH

For Arc-TRAP mice, three to four brain sections (20 µm thick) spaced 350–400 µm apart were analyzed per animal to span the dorsal striatum (from approximately +1.1 mm to −0.1 mm relative to bregma). ROIs covering dorsal striatum in both hemispheres were manually defined in ImageJ, and image analysis was performed using CellProfiler^44^ (v4.2.1; www.cellprofiler.org) with a custom-built pipeline. First, the “IdentifyPrimaryObjects” module was applied on DAPI channel to segment nuclei using adaptive thresholding and size filtering. Identified nuclei were then expanded by three pixels (∼ 1 micron) to generate cell body mask. Mean fluorescence intensities for Arc and tdTomato channels were measured for each cell. Thresholds for classifying Arc- and tdTomato-positive cells were determined using the “ClassifyObjects” and “FliterObjects” module on a representative subset of images and then applied uniformly across all sections. To identify double-positive cells, the “RelateObjects” module was used, with tdTomato-positive cells designated as “parent” objects and Arc-positive cells as “child” objects. A cell was classified as double positive when it was identified as related objects.

### Single-molecule RNA quantifications

To quantify DEGs in Fos-TRAP mice, a custom CellProfiler pipeline was optimized for single-molecule RNA ISH analysis. Two 20-µm-thick sections from primary motor cortex were processed and ROIs were manually selected and cropped in ImageJ. Nuclei were segmented in the DAPI channel using the “IdentifyPrimaryObjects” module, and a cell body mask was generated by expanding each nucleus by 8 pixels (∼1.2 µm). Mean fluorescence intensity in the tdTomato channel were measured, and TARP cells were identified using the “ClassifyObjects” and “FliterObjects” module. To detect individual RNA puncta within TRAP cells, the DEG channel was first enhanced using “EnhanceOrSuppressFeatures” module (Feature type: speckles) and then masked by the TRAP cell objects. RNA puncta were detected using the “IdentifyPrimaryObjects” module in the TRAP cells with a smaller object size range (1–10 pixels). Finally, RNA puncta in each TRAP cell were quantified using the “MaskObjects” and “RelateObjects” modules, with TRAP cells designated as both the mask and parent objects.

### scRNA-Seq data analysis

#### Generating scRNA expression matrices

scRNA expression matrices were generated by aligning sequencing reads to a custom reference using the Cell Ranger pipeline (v3.1.0. 10x Genomics). Briefly, a WPRE sequence (589 bp) located at 3’ end of Ai9 allele (Rosa-CAG-LSL-tdTomato-WPRE) was added at the end of mouse reference genome FASTA file and annotated in corresponding GTF file (*Mus musculus*, GRCm38/mm10, Ensemble database) to build a new reference (*cellranger mkref*). Base call files were demultiplexed into FASTAQ files (*cellranger mkfastq*) and aligned to the custom reference (*cellranger count*) following standard workflow of Cell Ranger. The output matrices were loaded into R (v4.2.3) for subsequent quality control, clustering and visualization.

#### Quality control, clustering and visualization

Seurat (v4) R package^45,46^ was used for the analysis of single-cell data using default parameters unless specified. All 10x expression matrices were loaded into Seurat and merged by their experimental groups. Data quality of each group was visualized by violin plots for nFeature_RNA, nCount_RNA and percent.mt (Figure S2A and S2B). Genes detected in fewer than 5 cells, and cells with fewer than 500 genes or more than 5% reads as mitochondrial were filtered out. We then split the data by their original matrices for the following normalization and integration (*SplitObject* function). Raw counts of each cell were normalized by SCTransform method ^47^ during which variations associated mitochondrial contamination were removed (*SCTransform* function, *vars.to.regress = “percent.mt“*). The top 3000 variable genes were selected to calculate Pearson residuals (*PrepSCTIntegration* function, *nfeatures = 3000*) and used to identify anchors (*FindIntegrationAnchors* function) for generating an integrated matrix (*IntegrateData* function). Next, principal component analysis (PCA) was applied to reduce the dimensions of integrated dataset (*RunPCA* function, *npcs = 50*). The first 50 significant PCs were used to construct a Shared Nearest Neighbor (SNN) Graph (*FindNeighbors* function, *dims = 1:50*) which was then used to determine cell clusters (*FindClusters* function, *resolution= 2.0*). To visualize the clusters, Uniform Manifold Approximation and Projection (UMAP) (*RunUMAP* function, *dims = 1:50*) and t-distributed stochastic neighbor embedding (t-SNE) (*RunTSNE* function, *dims = 1:50*) methods were performed for dimension reduction.

#### Cell type annotation

To identify the biological markers of each cell cluster, we performed differential gene expression tests on each cluster against all other clusters (*FindConservedMarkers* function, *grouping.var = “groups”*). The top 20 significantly enriched genes were considered as markers for each cluster. By comparing with previously known cell type markers, we grouped clusters expressing similar marker genes into major cell types, including glutamatergic neuron, GABAergic neuron, spiny projection neuron (SPN), oligodendrocyte (OL) & oligodendrocyte precursor cell (OPC), astrocyte (AS), microglia (MG), blood vessel (BV), neurogenesis and fibroblast_like cell. Clusters that co-expressed marker genes from more than two major cell types were classified as multiplex cell clusters and were excluded from further analysis.

Clusters of glutamatergic neurons and GABAergic neurons showed distinct marker gene expression patterns consistent with known neuronal subtypes. Therefore, we further classify them into subtypes. In motor cortex, we identified 8 glutamatergic projection neuron and 9 GABAergic interneuron subtypes: layer 2/3 intratelencephalic (L2/3 IT), L4/5 IT, L5/6 IT-1, 2, 3, L6 IT, L5 near projecting (L5 NP), L5 PT, L6 CT and L6b neurons, Parvalbumin-expressing interneurons (PV IN), Sst IN, Lamp5 IN, Enpp2 IN, Htr3a IN, Deptor IN, Pthlh IN, Crh IN and Crhbp IN. In striatum, we identified 3 SPN and 4 interneuron subtypes: direct pathway SPN (dSPN), indirect pathway SPN (iSPN), eccentric SPN (eSPN), Sst IN, PV IN, Th IN and Chat IN. DropViz^31^ (http://dropviz.org/) and Cell Type Knowledge Explorer^32^ (https://knowledge.brain-map.org/celltypes/CCN202002013) were used for selecting marker genes.

#### TRAP neurons identification

The expression of tdTomato (measured by counts mapping to WPRE) was used to identify neurons activated during behavior (TRAP-tdTom neurons). Specifically, all counts of neuronal clusters were normalized using counts per million (CPM) (*NormalizeData* function, *normalization.method = “RC”*, *scale.factor =1e6*). Cells with tdTomato counts > 10 CPM were determined as TRAP-tdTom neurons.

#### Differentially expressed gene (DEG) analysis

DEG analysis was performed on all genes (exclude mitochondrial genes) for TRAP neurons within each neuronal type and all cells within each glial type. Specifically, for each cell type, a Wilcoxon rank-sum test was applied on each gene, comparing expression level in cells from the early or late group to the ctrl group (*FindAllMarkers* function, *test.use = “wilcox”*), followed by Benjamini-Hochberg correction^48^ for multiple testing. Genes with an absolute log_2_ fold change > 0.26 and an adjusted p-value < 0.05 were considered as DEGs. To eliminate rare transcripts, genes detected in fewer than 10% of cells within a given cell type were excluded (*min.pct = 0.01*). In addition, gene expressions were compared between experimental batches and between TRAP and non-TRAP neurons in the ctrl group. Genes showing significant differences in these comparisons were removed from the DEG lists to minimize batch effects and exclude genes associated with general neuronal activation. DEGs were annotated using Metascape^49^ (https://metascape.org/) and visualized using volcano plots (EnhancedVolcano) and heatmaps (pheatmap).

#### Pathway enrichment analysis

Pathway enrichment analysis for DEGs was performed using the multigene list meta-analysis tool Metascape with default parameters. Briefly, a hypergeometric test was performed to compare each DEG list against predefined gene sets (gene memberships) associated with GO/KEGG terms, followed by Benjamini-Hochberg correction for multiple testing. All genes in the mouse genome were used as the enrichment background. Terms with a p-value < 0.01, gene count > 3, and an enrichment factor > 1.5 (ratio of observed counts to the counts expected by chance) were determined as significantly enriched terms. To reduce redundancy, Kappa similarity was calculated between enriched terms based on the overlap in their gene memberships. A hierarchical clustering tree was then constructed, and subtrees with a similarity score > 0.3 were defined as clusters. The most statistically significant term within each cluster was selected as its representative and was referred to as a DEGs-enriched pathway or cellular component in this paper.

### Behavior tracking and analysis

The head-fixed forelimb pellet-reaching task was recorded at 100 fps using an infrared camera and analyzed with DeepLabCut^50^ (DLC, v2.1.0). To create a DLC network, 433 frames were automatically extracted from videos across six animals using k-means clustering to generate training datasets. Key body parts were manually labeled in each frame, including the nose, eye, food pellet, dominant paw, non-dominant paw and tongue. The DLC network was trained on 95% of the datasets using MobileNet v2.1.0-based architecture for 1,030,000 iterations. Model performance was evaluated using the remaining 5% of the datasets by computing the root mean square error (RMSE) between predicted and manually annotated points. After training, the test error with p-cutoff is 3.62 pixels. Then, all video frames were analyzed using the trained DLC network to extract body parts positions (*x-y* coordinates in pixels) for each frame. Low-confidence detections (likelihood < 0.9) were excluded, and x-y trajectories were smoothened using a 2-point running mean with a custom MATLAB script. Dominant paw velocities were calculated using sqrt (Δx^2^ + Δy^2^) / Δt.

All videos were manually inspected frame-by-frame using BORIS (Behavioral Observation Research Interactive Software)^51^ and annotated for key behaviors, including reaching (success, failed and in-vain) and chewing. The start of a reaching event was defined as the frame preceding a paw movement greater than 10 pixels, and the end frame was marked when the paw contacted either the pellet or the dispenser. Chewing events were annotated from the moment the pellet entered the mouth until mouth movement ceased. All labeled behavioral data were exported in CSV format and used to synchronize Ca²⁺ imaging data with behavior and extract reaching bouts *via* a custom MATLAB script.

### Analysis of *in vivo* two-photon imaging data

All recorded two-photon fluorescence time-lapse images were first corrected for any DC offset in the pixel values. Therefore, the minimum pixel value of each image frame was averaged across all frames. This average minimum pixel value was then subtracted from all pixels in the time-lapse image. Next, images were corrected for slow changes in image brightness over the course of each imaging session, such as caused by mild bleaching of GCaMP. For this, the mean pixel value of the first image frame was extracted and a linear fit of the brightness as a function of each frame was calculated. Each frame was then divided by its fitted brightness. Lastly, images were corrected for brain motion during imaging with a piecewise non-rigid motion correction approach (NoRMCorre)^52^ using 256 X 256 pixel patches of each image. To extract Ca^2+^ traces from individual neurons, ROIs in ImageJ/FIJI were manually selected and the average pixel values were extracted across time.

### Statistics

Statistical analyses were performed using R for scRNA-seq data and RNA ISH validations, Prism 9.0 (GraphPad Software) for brain histology and behavior, and MATLAB for calcium imaging. Statistics tests were described in Figure legends. All error bars represent standard error of mean, SEM.

## Supplemental information

**Supplemental Figure S1-S7**

**Table S1.** List of DEGs in each neuronal subtype, related to Figure 4 and S5.

Tab 1: Differentially expressed genes (DEGs) between Early TRAP and Ctrl TRAP for each neuronal subtype in the motor cortex.

Tab 2: DEGs between Late TRAP and Ctrl TRAP for each neuronal subtype in the motor cortex.

Tab 3: DEGs between Early TRAP and Ctrl TRAP for each neuronal subtype in the striatum.

Tab 4: DEGs between Late TRAP and Ctrl TRAP for each neuronal subtype in the striatum.

In each tab, columns (from left to right) include: Gene Name, raw p-value from the Wilcoxon rank-sum test, adjusted p-value using the Holm-Bonferroni correction, average log₂ fold change, proportion of cells expressing the gene in Early or Late TRAP, proportion of cells expressing the gene in Ctrl TRAP, Cell Type, and Comparison group. Only genes with an adjusted p-value < 0.05 and an absolute log₂ fold change > 0.26 are included.

**Table S2.** List of DEGs in each glial subtype, related to Figure 6 and S7.

Tab 1: DEGs between Early and Ctrl for each glia subtype in the motor cortex. Tab 2: DEGs between Late and Ctrl for each glia subtype in the motor cortex. Tab 3: DEGs between Early and Ctrl for each glia subtype in the striatum.

Tab 4: DEGs between Late and Ctrl for each glia subtype in the striatum.

In each tab, columns (from left to right) include: Gene Name, raw p-value from the Wilcoxon rank-sum test, adjusted p-value using the Holm-Bonferroni correction, average log₂ fold change, proportion of cells expressing the gene in Early or Late, proportion of cells expressing the gene in Ctrl , Cell Type, and Comparison group. Only genes with an adjusted p-value < 0.05 and an absolute log₂ fold change > 0.26 are included.

