## Supplemental Figures for "Motor learning drives region-specific transcriptomic remodeling in the motor cortex and dorsal striatum"

Figure S1

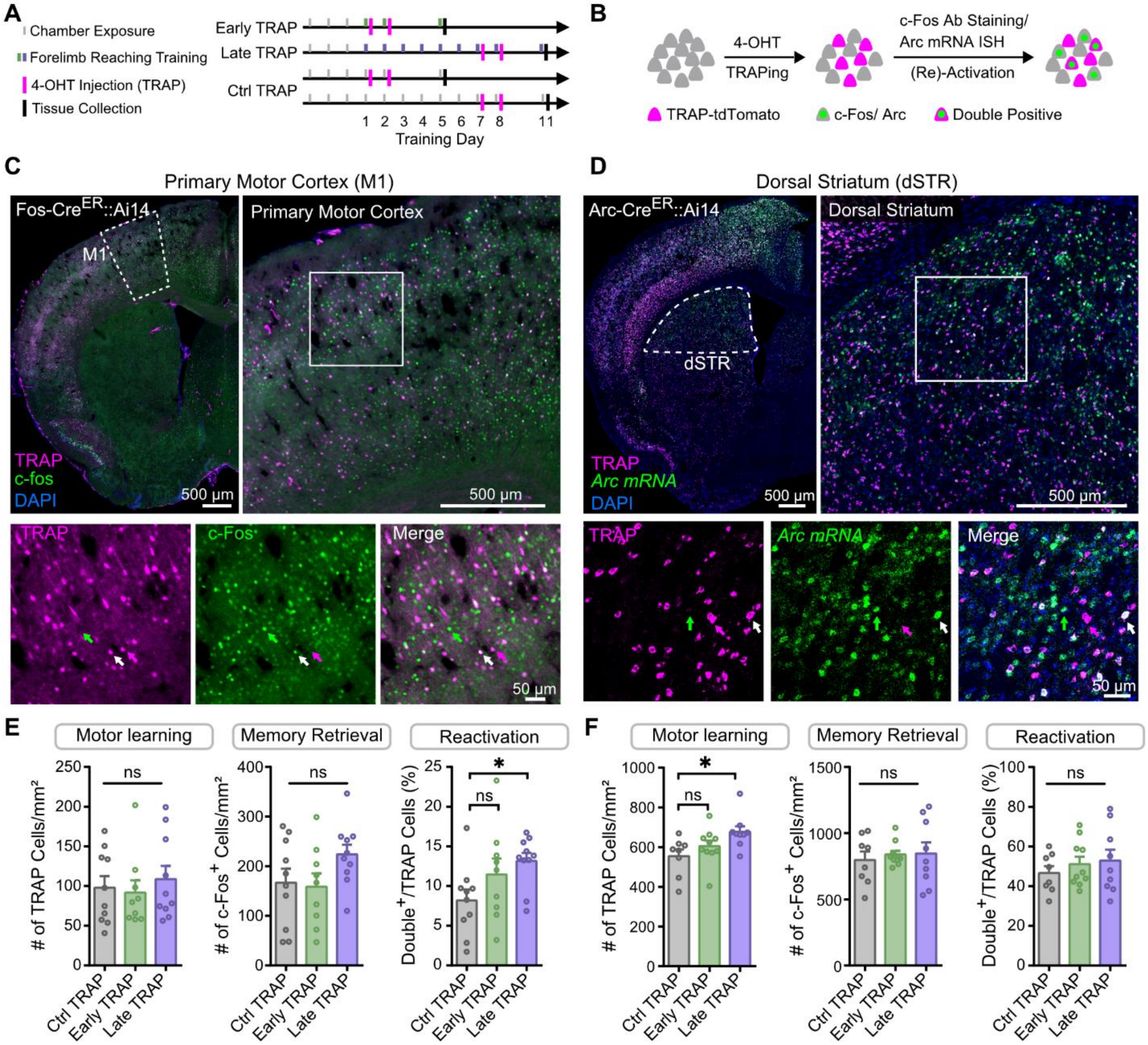

A
Chamber Exposure
Forelimb Reaching Training
4-OHT Injection (TRAP)
Tissue Collection
Early TRAP
Late TRAP
Ctrl TRAP
1 2 3 4 5 6 7 8 11
Training Day
B
4-OHT
TRAPing
c-Fos Ab Staining/
Arc mRNA ISH
(Re)-Activation
TRAP-tdTomato
c-Fos/ Arc
Double Positive
C
Primary Motor Cortex (M1)
Fos-CreER::Ai14
M1
Primary Motor Cortex
TRAP
c-fos
DAPI
500 µm
500 µm
TRAP
c-Fos
Merge
50 µm
D
Dorsal Striatum (dSTR)
Arc-CreER::Ai14
dSTR
Dorsal Striatum
TRAP
Arc mRNA
DAPI
500 µm
500 µm
TRAP
Arc mRNA
Merge
50 µm
E
Motor learning
Memory Retrieval
Reactivation
### of TRAP Cells/mm²
### of c-Fos+ Cells/mm²
Double+/TRAP Cells (%)
ns
ns
*
ns
Ctrl TRAP
Early TRAP
Late TRAP
F
Motor learning
Memory Retrieval
Reactivation
### of TRAP Cells/mm²
### of c-Fos+ Cells/mm²
Double+/TRAP Cells (%)
*
ns
ns
ns
Ctrl TRAP
Early TRAP
Late TRAP

**Figure S1. TRAP-labeled neurons are behaviorally relevant during motor learning and memory retrieval, related to Figure 1**

(A) Experimental timeline of the reaching task and TRAP labeling during motor learning and memory retrieval phases.

(B) Schematic of the experimental design used to identify activated neuronal populations during learning (TRAP-tdTomato) and memory retrieval (via c-fos immunostaining or *Arc* RNA in situ hybridization [ISH]).

(C) Representative images of TRAP-tdTomato+ cells (magenta) and c-fos+ cells (green) in the primary motor cortex (M1). Top left: Overview of a brain hemisphere. Top right: Enlarged view of M1. Bottom: Magnified view of the solid square in M1. Green arrows indicate c-fos+ only neurons; magenta arrows, TRAP-labeled neurons only; white arrows, neurons co-expressing c-fos and TRAP-tdTomato.

(D) Representative images of TRAP-tdTomato+ cells (magenta) and Arc RNA ISH signal (green) in the dorsal striatum (dSTR). Top left: hemisphere overview. Top right: Enlarged view of dSTR. Bottom: Magnified region indicated by the solid square in dSTR. Arrows follow the same labeling conventions as in (C).

(E) Quantification of TRAP-tdTomato+ cells (left), c-fos+ cells (middle) and proportion of TRAP and c-fos double positive cells (right) in Ctrl-TRAP (n= 10), Early-TRAP (n= 9) and Late-TRAP (n= 10) mice. Bars represent mean values; individual data points represent individual mice. Error bars, SEM. One-way ANOVA with Tukey's multiple comparison test. $^*p < 0.05$; ns, non-significant.

(F) Quantification of TRAP-tdTomato+ cells (left), *Arc*+ cells (middle) and proportion of TRAP and *Arc* double positive cells (right) in Ctrl-TRAP (n= 8), Early-TRAP (n= 10) and Late-TRAP (n= 9) mice. Data are shown as in (E). Error bars, SEM. One-way ANOVA with Tukey's multiple comparison test. $^*p < 0.05$; ns, non-significant.

Figure S2

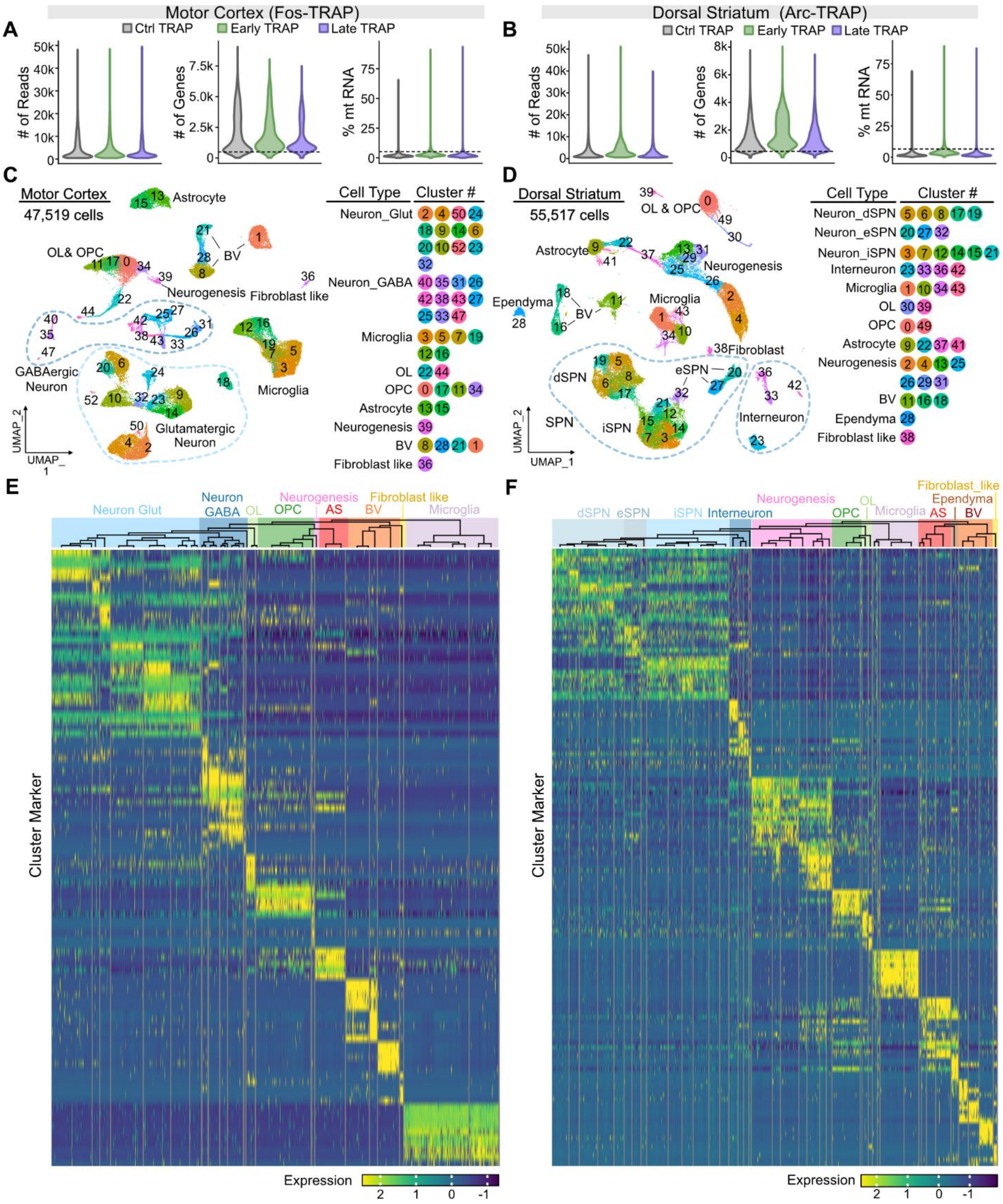

Motor Cortex (Fos-TRAP)
Dorsal Striatum (Arc-TRAP)
Ctrl TRAP
Early TRAP
Late TRAP
### of Reads
### of Genes
% mt RNA
Motor Cortex
47,519 cells
Astrocyte
OL& OPC
BV
Neurogenesis
Fibroblast like
GABAergic Neuron
Glutamatergic Neuron
Microglia
UMAP_2
UMAP_1
Cell Type
Cluster #
Neuron_Glut
Neuron_GABA
Microglia
OL
OPC
Astrocyte
Neurogenesis
BV
Fibroblast like
Dorsal Striatum
55,517 cells
OL & OPC
Astrocyte
Neurogenesis
Ependyma
Microglia
Fibroblast
dSPN
eSPN
iSPN
SPN
Interneuron
Neuron_dSPN
Neuron_eSPN
Neuron_iSPN
Interneuron
Microglia
OL
OPC
Astrocyte
Neurogenesis
BV
Ependyma
Fibroblast like
Neuron Glut
Neuron GABA
OL
OPC
Neurogenesis
AS
BV
Fibroblast like
Microglia
dSPN
eSPN
iSPN
Interneuron
Neurogenesis
OPC
OL
Microglia
AS
Ependyma
BV
Fibroblast_like
Cluster Marker
Expression

**Figure S2. Quality control, clustering, and cell type annotations of scRNA-seq data from the motor cortex and dorsal striatum, related to Figure 1**

(A) Violin plots showing the number of reads (left), number of genes (middle), and percentage of mitochondrial RNA (right) of all cells recovered from the motor cortex. Dashed lines indicate quality control thresholds: <500 genes or >5% mitochondrial RNA content.

(B) Violin plots showing the number of reads (left), number of genes (middle), and percentage of mitochondrial RNA (right) of all cells recovered from the dorsal striatum. Dashed lines represent the same filtering thresholds as in (A).

(C) UMAP visualization of 52 clusters from the motor cortex classified into major neuronal and glial cell types. Colors represent individual clusters.

(D) UMAP visualization of 49 clusters in the dorsal striatum classified into major neuronal and glial cell types. Colors represent individual clusters.

(E) Marker genes-based cell type annotation in the motor cortex. Top: A dendrogram illustrating the relationships among clusters based on transcriptome similarity. Colors indicate major neuronal and glial cell types. Bottom: Heatmap of the top five marker genes for each cluster. Columns correspond to individual cells, rows represent genes, ordered by clusters.

(F) Marker genes-based cell type annotation in the dorsal striatum. Top: A dendrogram of transcriptomic cluster relationships; Bottom: Heatmap of the top five marker genes for each cluster. Colors and format as in (E).

Abbreviations: OL, oligodendrocyte; OPC, oligodendrocyte precursor cells; BV, blood vessels; SPN, spiny projection neuron.

Figure S3

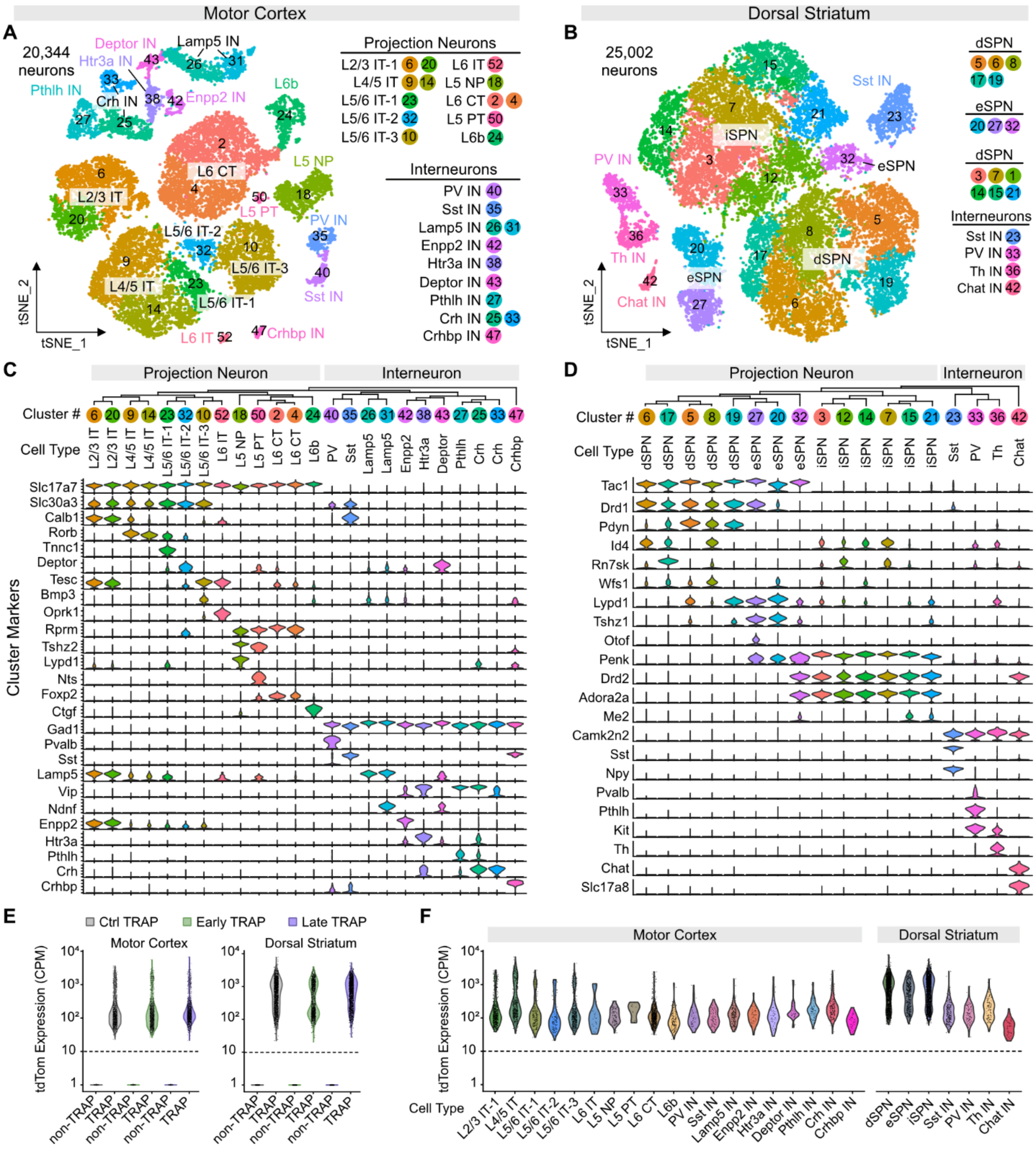

Motor Cortex
Dorsal Striatum
A
20,344 neurons
Projection Neurons
L2/3 IT-1 6 20
L4/5 IT 9 14
L5/6 IT-1 23
L5/6 IT-2 32
L5/6 IT-3 10
L6 IT 52
L5 NP 18
L6 CT 2 4
L5 PT 50
L6b 24
Interneurons
PV IN 40
Sst IN 35
Lamp5 IN 26 31
Enpp2 IN 42
Htr3a IN 38
Deptor IN 43
Pthlh IN 27
Crh IN 25 33
Crhbp IN 47
tSNE_1
tSNE_2
B
25,002 neurons
dSPN 5 6 8 17 19
eSPN 20 27 32
dSPN 3 7 1 14 15 21
Interneurons
Sst IN 23
PV IN 33
Th IN 36
Chat IN 42
C
Projection Neuron
Interneuron
Cluster #
Cell Type
Cluster Markers
Slc17a7
Slc30a3
Calb1
Rorb
Tnnc1
Deptor
Tesc
Bmp3
Oprk1
Rprm
Tshz2
Lypd1
Nts
Foxp2
Ctgf
Gad1
Pvalb
Sst
Lamp5
Vip
Ndnf
Enpp2
Htr3a
Pthlh
Crh
Crhbp
D
Projection Neuron
Interneuron
Cluster #
Cell Type
Tac1
Drd1
Pdyn
Id4
Rn7sk
Wfs1
Lypd1
Tshz1
Otof
Penk
Drd2
Adora2a
Me2
Camk2n2
Sst
Npy
Pvalb
Pthlh
Kit
Th
Chat
Slc17a8
E
Ctrl TRAP
Early TRAP
Late TRAP
Motor Cortex
Dorsal Striatum
tdTom Expression (CPM)
non-TRAP
TRAP
F
Motor Cortex
Dorsal Striatum
tdTom Expression (CPM)
Cell Type
L2/3 IT-1
L4/5 IT
L5/6 IT-1
L5/6 IT-2
L5/6 IT-3
L6 IT
L5 NP
L5 PT
L6 CT
L6b
PV IN
Sst IN
Lamp5 IN
Enpp2 IN
Htr3a IN
Deptor IN
Pthlh IN
Crh IN
Crhbp IN
dSPN
eSPN
iSPN
Sst IN
PV IN
Th IN
Chat IN

**Figure S3. Classification of neuronal clusters and identification of TRAP-labeled neurons, related to Figure 2**

(A) t-SNE plot showing 24 neuronal clusters from the motor cortex classified into 19 neuronal subtypes, color-coded by cluster identity.

(B) t-SNE plot showing 18 neuronal clusters from the dorsal striatum classified into 7 neuronal subtypes, color-coded by cluster identity.

(C) Marker genes used to classify neuronal subtypes in the motor cortex. Top: Dendrogram illustrating the transcriptional similarity among clusters. Bottom: Violin plots showing the expression levels of representative marker genes for each subtype.

(D) Marker genes used to classify neuronal subtypes in the dorsal striatum. Top: Dendrogram illustrating the transcriptional similarity among clusters. Bottom: Violin plots showing the expression levels of representative marker genes for each subtype.

(E) Violin plots showing tdTomato expression levels in TRAP and non-TRAP neurons across Ctrl, Early and Late TRAP groups in the motor cortex (left) and dorsal striatum (right). Dashed lines indicate the threshold for defining TRAP neurons: >10 CPM (counts per million) of tdTomato reads per cell.

(F) Violin plots showing tdTomato expression in TRAP neurons across neuronal subtypes in the motor cortex (left) and dorsal striatum (right). Dashed lines mark the TRAP cell threshold (>10 CPM).

Abbreviations: IT, intratelencephalically projecting; NP, near-projecting; PT, pyramidal tract; CT, corticothalamic projecting; PV, parvalbumin; SPN, spiny projection neuron.

Figure S4

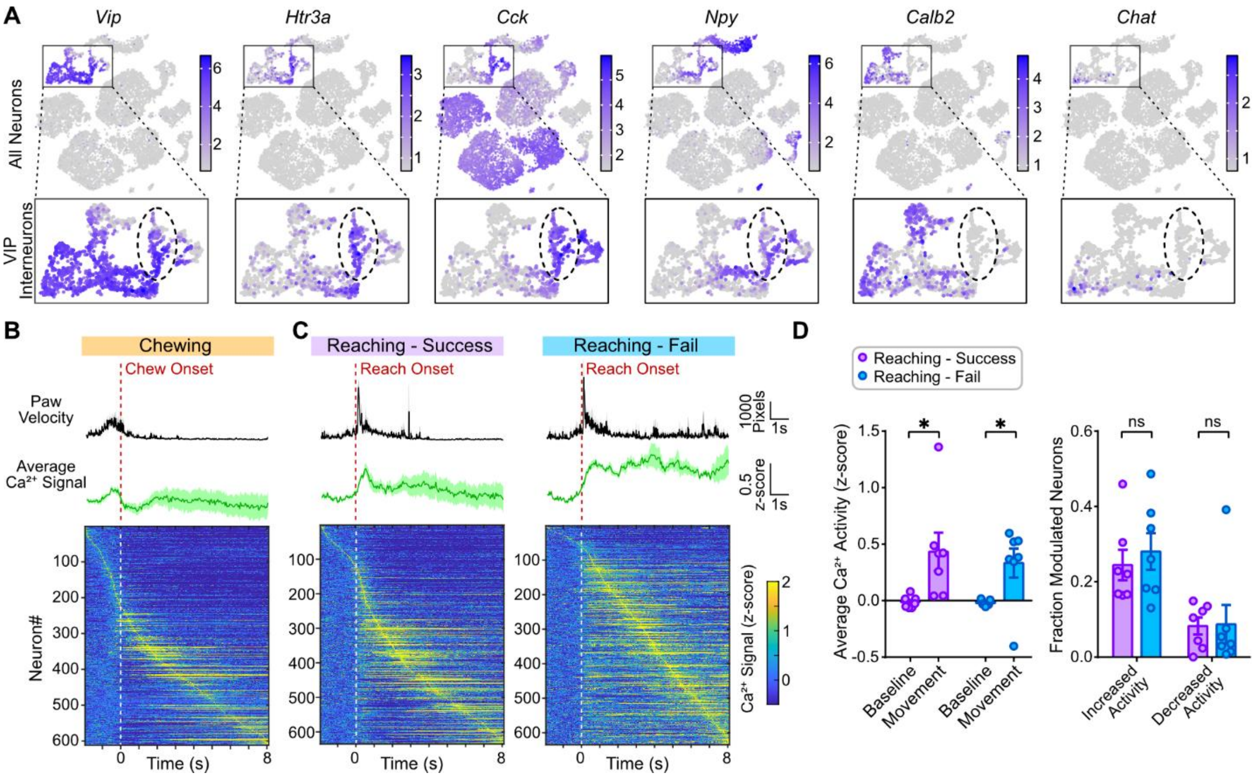


**Figure S4. Htr3a-expressing interneurons are specifically activated during learned reaching behavior, related to Figure 3**

(A) Feature plots showing expression of canonical VIP interneuron maker genes. Insets show the distribution of four VIP interneuron subtypes.

(B) $Ca^{2+}$ activity of Htr3a interneurons during chewing. Top: Average paw velocity across all chewing bouts from 6 mice (mean ± SEM). Middle: Average $Ca^{2+}$ signals of Htr3a neurons during chewing (mean ± SEM). Bottom: Heatmap of $Ca^{2+}$ activity from all Htr3a neurons ordered by peak activation time. Each row represents one individual neuron. Dashed line indicates onset of chewing behavior.

(C) $Ca^{2+}$ activity of Htr3a neurons during successful (left) and failed (right) reaching. Top: Average paw movement velocity for successful (left) or failed (right) reaching bouts from (mean ± SEM, n=7 mice). Middle: Average $Ca^{2+}$ signals of all Htr3a neurons during successful (left) or failed (right) reaching (mean ± SEM). Bottom: Heatmap showing individual neuronal $Ca^{2+}$ signals ordered by peak activation time. Dashed line indicates reach onset.

(D) Left: average $Ca^{2+}$ signals at baseline and during reaching (left), separated into successful and failed reaches. Right: fraction of Htr3a neurons modulated during each condition. Bars represent group means; dots indicate individual mice (*n* = 7). Error bars, SEM. n = 7 mice. Wilcoxon matched pairs signed rank test. *p < 0.05; ns, non-significant.

Figure S5

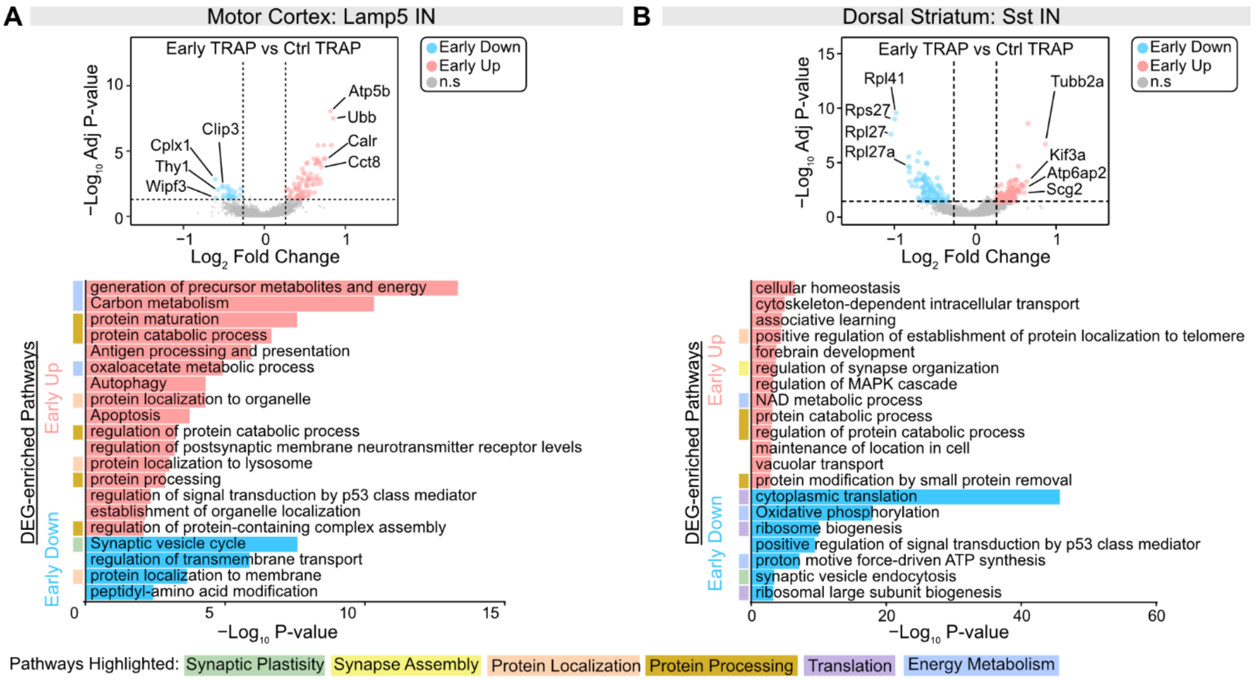


**Figure S5. DEG and pathway analysis for interneurons, related to Figure 4 and 5.**

(A) DEG and pathway analysis in cortical Lamp5 interneurons. Top: volcano plot showing $\log_2$ fold change and $-\log_{10}$ adjusted *p*-values for all genes comparing Early TRAP vs. Ctrl TRAP. Wilcox rank sum test followed by Holm-Bonferroni correction. ns, non-significant. DEG Cutoff: Adjusted p-value < 0.05 and |Log2 Fold Change| > 0.26. Bottom: Bar plots of GO and KEGG pathway enrichment for upregulated and downregulated DEGs in Early TRAP Lamp5 INs. Hypergeometric test with Benjamini-Hochberg correction. Significant terms (p < 0.01, gene count > 3, enrichment factor > 1.5) were clustered by Kappa similarity (> 0.3), and the top 20 enriched pathways are shown. Shaded boxes group pathways by biological classifications of each term (as indicated in the list at bottom).

(B) DEG and pathway enrichment analysis in striatal Sst interneurons. Top: volcano plot comparing Early TRAP vs. Ctrl TRAP gene expression in Sst INs, analyzed as in (A). Bottom: Bar plots showing enriched GO and KEGG pathways among upregulated and downregulated DEGs in Early TRAP Sst interneurons, analyzed as in (A). Biological classifications are indicated by shaded boxes (list at bottom).

Figure S6

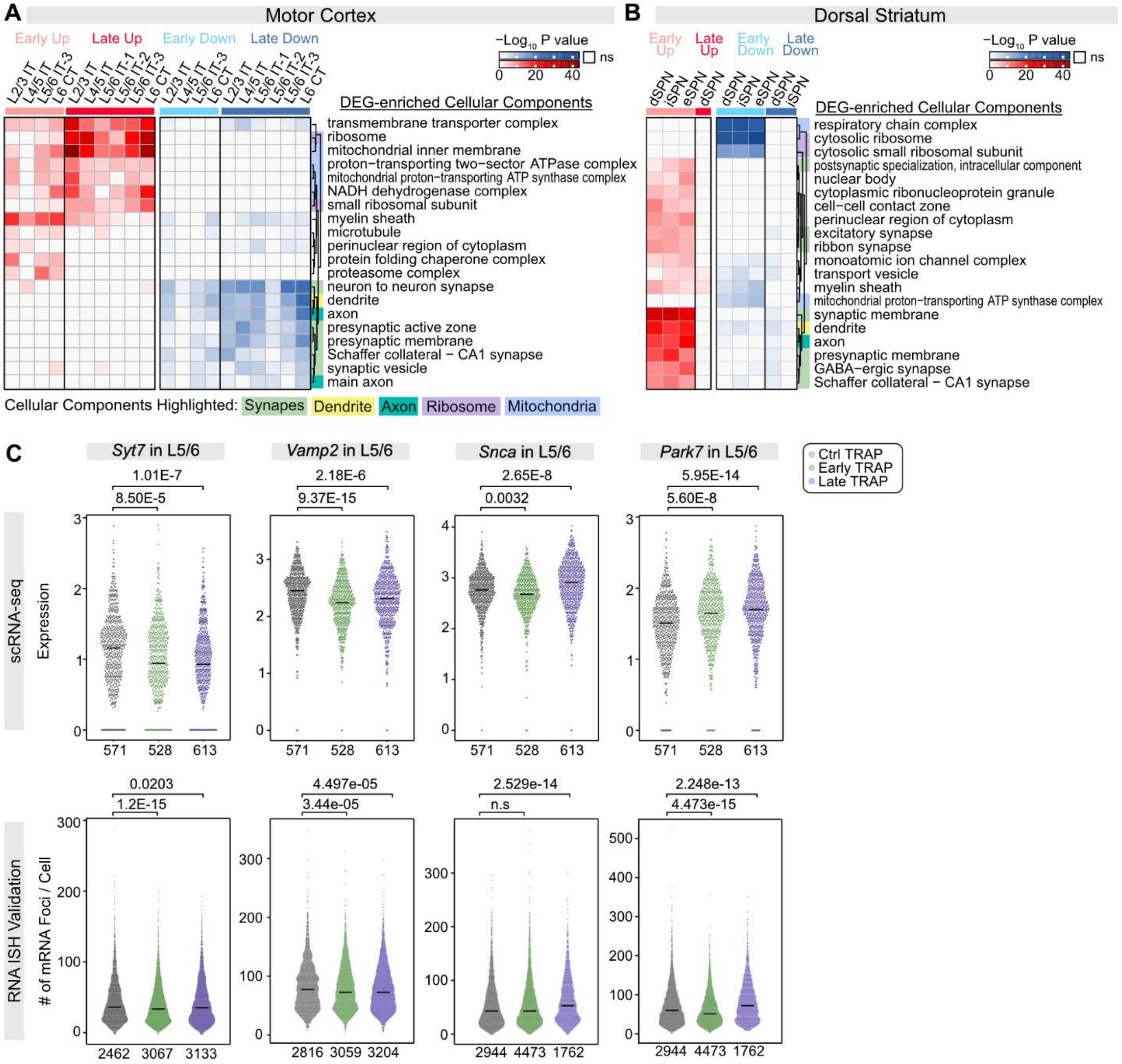
A
Motor Cortex
Early Up
Late Up
Early Down
Late Down
−Log10 P value
ns
0 20 40
L2/3 IT
L4/5 IT
L5/6 IT-3
L6 CT
L2/3 IT
L4/5 IT
L5/6 IT-1
L5/6 IT-2
L5/6 IT-3
L6 CT
DEG-enriched Cellular Components
transmembrane transporter complex
ribosome
mitochondrial inner membrane
proton−transporting two−sector ATPase complex
mitochondrial proton−transporting ATP synthase complex
NADH dehydrogenase complex
small ribosomal subunit
myelin sheath
microtubule
perinuclear region of cytoplasm
protein folding chaperone complex
proteasome complex
neuron to neuron synapse
dendrite
axon
presynaptic active zone
presynaptic membrane
Schaffer collateral − CA1 synapse
synaptic vesicle
main axon
Cellular Components Highlighted: Synapses Dendrite Axon Ribosome Mitochondria
B
Dorsal Striatum
Early Up
Late Up
Early Down
Late Down
−Log10 P value
ns
0 20 40
dSPN
iSPN
eSPN
DEG-enriched Cellular Components
respiratory chain complex
cytosolic ribosome
cytosolic small ribosomal subunit
postsynaptic specialization, intracellular component
nuclear body
cytoplasmic ribonucleoprotein granule
cell−cell contact zone
perinuclear region of cytoplasm
excitatory synapse
ribbon synapse
monoatomic ion channel complex
transport vesicle
myelin sheath
mitochondrial proton−transporting ATP synthase complex
synaptic membrane
dendrite
axon
presynaptic membrane
GABA−ergic synapse
Schaffer collateral − CA1 synapse
C
Syt7 in L5/6
Vamp2 in L5/6
Snca in L5/6
Park7 in L5/6
Ctrl TRAP
Early TRAP
Late TRAP
scRNA-seq
Expression
1.01E-7
8.50E-5
2.18E-6
9.37E-15
2.65E-8
0.0032
5.95E-14
5.60E-8
571 528 613
RNA ISH Validation
### of mRNA Foci / Cell
0.0203
1.2E-15
2462 3067 3133
4.497e-05
3.44e-05
2816 3059 3204
2.529e-14
n.s
2944 4473 1762
2.248e-13
4.473e-15
2944 4473 1762

**Figure S6. Enrichment of cellular components in projection neurons and *in situ* hybridization (ISH) validation of key DEGs, related to Figure 5.**

(A) Cellular components enrichment analysis of DEGs in projection neuron subtypes from the motor cortex. Heatmap showing $-\log_{10}$ P values with red indicating upregulated and blue indicating downregulated pathways. Hypergeometric test with Benjamini-Hochberg correction. Significant terms ($p < 0.01$, gene count > 3, enrichment factor> 1.5) were clustered by Kappa similarity (> 0.3). Rows represent the top 20 enriched terms; columns represent subtypes. Term names are displayed in a dendrogram (right) with shaded boxes denoting their cellular locations (list at bottom).

(B) Cellular components enrichment analysis of DEGs in projection neuron subtypes from the dorsal striatum. Heatmap formatting, organization and statistics as in (A).

(C) Scatter violin plots showing the expression levels of DEGs in Ctrl, Early, and Late TRAP cells. Top: expression based on scRNA-seq data. Bottom: expression validated by signal-molecule RNA ISH. Scatter points represent cells. Black crossbars indicate group medians. Wilcox rank sum test. P values are displayed above each violin plot; cell numbers are shown below. ns, non-significant.

Abbreviations: IT, intratelencephalically projecting; NP, near-projecting; PT, pyramidal tract; CT, corticothalamic projecting; PV, parvalbumin; SPN, spiny projection neuron.

Figure S7

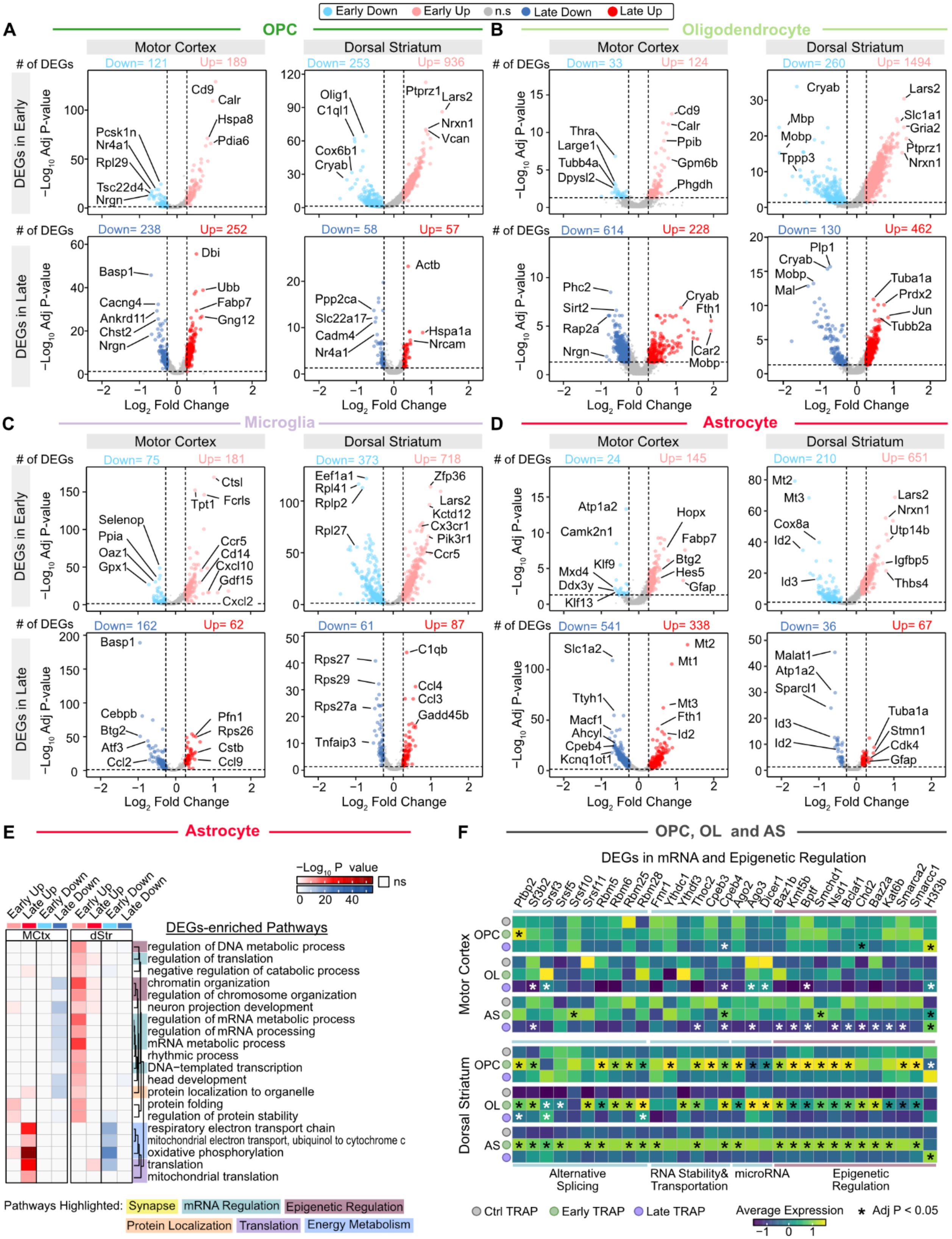

Early Down
Early Up
n.s
Late Down
Late Up
A
OPC
B
Oligodendrocyte
Motor Cortex
Dorsal Striatum
### of DEGs
Down= 121
Up= 189
Down= 253
Up= 936
Down= 33
Up= 124
Down= 260
Up= 1494
DEGs in Early
DEGs in Late
−Log10 Adj P-value
Cd9
Calr
Hspa8
Pdia6
Pcsk1n
Nr4a1
Rpl29
Tsc22d4
Nrgn
Olig1
C1ql1
Cox6b1
Cryab
Ptprz1
Lars2
Nrxn1
Vcan
Thra
Large1
Tubb4a
Dpysl2
Ppib
Gpm6b
Phgdh
Mbp
Mobp
Tppp3
Slc1a1
Gria2
Down= 238
Up= 252
Down= 58
Up= 57
Down= 614
Up= 228
Down= 130
Up= 462
Dbi
Basp1
Ubb
Fabp7
Gng12
Cacng4
Ankrd11
Chst2
Actb
Ppp2ca
Slc22a17
Cadm4
Nr4a1
Hspa1a
Nrcam
Phc2
Sirt2
Rap2a
Fth1
Car2
Plp1
Mal
Tuba1a
Prdx2
Jun
Tubb2a
Log2 Fold Change
C
Microglia
D
Astrocyte
Down= 75
Up= 181
Down= 373
Up= 718
Down= 24
Up= 145
Down= 210
Up= 651
Ctsl
Tpt1
Fcrls
Selenop
Ppia
Oaz1
Gpx1
Ccr5
Cd14
Cxcl10
Gdf15
Cxcl2
Eef1a1
Rpl41
Rplp2
Rpl27
Zfp36
Kctd12
Cx3cr1
Pik3r1
Atp1a2
Camk2n1
Klf9
Mxd4
Ddx3y
Klf13
Hopx
Btg2
Hes5
Gfap
Mt2
Mt3
Cox8a
Id2
Id3
Utp14b
Igfbp5
Thbs4
Down= 162
Up= 62
Down= 61
Up= 87
Down= 541
Up= 338
Down= 36
Up= 67
Cebpb
Atf3
Ccl2
Pfn1
Rps26
Cstb
Ccl9
Rps27
Rps29
Rps27a
Tnfaip3
C1qb
Ccl4
Ccl3
Gadd45b
Slc1a2
Ttyh1
Macf1
Ahcyl
Cpeb4
Kcnq1ot1
Mt1
Malat1
Sparcl1
Stmn1
Cdk4
E
Astrocyte
−Log10 P value
ns
Early Up
Late Up
Early Down
Late Down
MCtx
dStr
DEGs-enriched Pathways
regulation of DNA metabolic process
regulation of translation
negative regulation of catabolic process
chromatin organization
regulation of chromosome organization
neuron projection development
regulation of mRNA metabolic process
regulation of mRNA processing
mRNA metabolic process
rhythmic process
DNA−templated transcription
head development
protein localization to organelle
protein folding
regulation of protein stability
respiratory electron transport chain
mitochondrial electron transport, ubiquinol to cytochrome c
oxidative phosphorylation
translation
mitochondrial translation
Pathways Highlighted: Synapse
mRNA Regulation
Epigenetic Regulation
Protein Localization
Translation
Energy Metabolism
F
OPC, OL and AS
DEGs in mRNA and Epigenetic Regulation
Ptbp2 Sf3b2 Srsf3 Srsf5 Srsf10 Srsf11 Rbm6 Rbm25 Rbm28 Fmr1 Ythdc1 Ythdf3 Thoc2 Cpeb3 Cpeb4 Ago2 Ago3 Dicer1 Baz1b Kmt5b Bptf Smchd1 Nsd1 Bclaf1 Chd2 Baz2a Kat6b Smarca2 Smarcc1 H3f3b
Motor Cortex
Dorsal Striatum
OPC
OL
AS
Alternative Splicing
RNA Stability& Transportation
microRNA
Epigenetic Regulation
Ctrl TRAP
Early TRAP
Late TRAP
Average Expression
Adj P < 0.05

**Figure S7. Identify motor learning-associated DEGs and functional pathways in glia cells, related to Figure 6.**

(A) Volcano plots showing all genes in oligodendrocyte precursor cell (OPC). Top row: Early vs Ctrl; Bottom row: Late vs Ctrl. Left column: motor cortex. right column: dorsal striatum. Wilcox rank sum test followed by Holm-Bonferroni correction. DEG Cutoff: |Log2 Fold Change| > 0.26 and Adjusted p-value < 0.05. ns, non-significant.

(B) Volcano plots showing all genes in oligodendrocyte (OL). Layout and statistics as in (A).

(C) Volcano plots showing all genes in microglia. Layout and statistics as in (A).

(D) Volcano plots showing all genes in astrocyte. Layout and statistics as in (A).

(E) GO and KEGG pathway enrichment analysis of DEGs in astrocytes. Heatmap displaying -$\log_{10}$ P values with red indicating upregulated pathways and blue indicating downregulated pathways. Hypergeometric test with Benjamini-Hochberg correction. Significant terms (p < 0.01, gene count > 3, enrichment factor > 1.5) were clustered by Kappa similarity (> 0.3), and the top 20 enriched pathways are shown. Term names are displayed in a dendrogram (right) with shaded boxes denoting their biological classifications (list at bottom).

(F) Expression of DEGs related to mRNA and epigenetic regulation in OPCs, OLs and astrocytes. Heatmaps showing average expression across Ctrl, Early, and Late TRAP neurons for each glial type of the motor cortex (Top) and dorsal striatum (bottom). DEG categories are labeled beneath the heatmaps. Wilcox rank sum test followed by Holm-Bonferroni correction. *Adjust p value < 0.05.

Abbreviations: OPC, oligodendrocyte precursor cell; OL, oligodendrocyte; AS, astrocyte; MG, microglia; MCtx, motor cortex; dSTR, dorsal striatum.
